# *ETV6* Deficiency and Microsatellite Enhancers Drive Transcriptional Dysregulation in B-Lymphoblastic Leukemia

**DOI:** 10.1101/2022.09.20.508499

**Authors:** Rohan Kodgule, Joshua W. Goldman, Alexander C. Monovich, Travis Saari, Cody N. Hall, Niharika Rajesh, Juhi Gupta, Athalee Aguilar, Noah A. Brown, Mark Y. Chiang, Marcin P. Cieslik, Russell J.H. Ryan

## Abstract

Distal enhancers play critical roles in sustaining oncogenic gene expression programs. We identify aberrant enhancer-like activation of GGAA tandem repeats as a characteristic feature of B-cell acute lymphoblastic leukemia (B-ALL) with genetic defects of the *ETV6* transcriptional repressor, including *ETV6-RUNX1*+ and *ETV6*-null B-ALL. We show that GGAA repeat enhancers are direct activators of previously identified *ETV6-RUNX1*+ B-ALL “signature” genes, including likely oncogenic drivers. When restored to ETV6-deficient B-ALL cells, ETV6 directly binds to GGAA repeat enhancers, represses their acetylation, downregulates adjacent genes, and inhibits B-ALL growth. In ETV6-deficient B-ALL cells, we find that the ETS transcription factor ERG directly binds to GGAA microsatellite enhancers and is required for sustained activation of many repeat enhancer-activated genes. Together, our findings reveal a novel epigenetic gatekeeper function of the *ETV6* tumor suppressor gene and establish microsatellite enhancers as a key mechanism underlying the unique gene expression program of *ETV6-RUNX1*+ B-ALL.

**Significance:** We show that the oncogenic gene expression program of a common pediatric leukemia relies on repetitive noncoding elements that are not conserved between humans and rodents, placing important limitations on animal models for this disease. Our findings may present new opportunities for targeting cancer-specific chromatin dysregulation in leukemia.

## Introduction

Integrative profiling of B-lymphoblastic leukemia (B-ALL) has identified clinically and biologically distinct subtypes associated with specific genomic alterations and gene expression signatures^1–3^. The *ETV6-RUNX1* (E-R) fusion oncogene defines a common B-ALL subtype, representing about 25% of pediatric B-ALL^4^. The encoded oncoprotein incorporates a protein-protein interaction domain of ETV6, a transcriptional repressor in the ETS family^5^, and nearly the full length of the DNA sequence-specific transcription factor (TF) RUNX1^6, 7^. Somatic mutations or genomic deletions that inactivate *ETV6* are frequently seen as secondary events in E-R*+* B-ALL^8–10^, in B-ALL cases that lack the E-R fusion gene but show an “*ETV6-RUNX1*-like” signature of aberrant gene expression^2, 11^, and as second-hit events in leukemias that arise in patients with germline loss-of-function *ETV6* variants^12, 13^. However, the mechanism by which ETV6 dysfunction contributes to leukemia is poorly understood. Here, we identify enhancer-like chromatin state and function of GGAA microsatellite repeats as a characteristic feature of *E-R*+ and *ETV6*-null B-ALL. Leukemia-specific GGAA repeat enhancers are bound by the ETS family activator ERG, sustain the expression of known E-R+ B-ALL signature genes, including potential leukemia drivers, and are repressed upon restoration of ETV6 expression. Our findings reveal an unexpected mechanism underlying the biology of a common subtype of childhood leukemia.

## Results

To identify novel features of enhancer dysregulation in B-cell malignancies, we performed genome-wide clustering of histone H3 lysine 27 acetylation (H3K27ac) ChIP-Seq and ATAC-Seq data from 26 B-cell cancer cell lines, including 13 B-ALL cell lines (Figure 1A-B, Suppl. Fig. 1A-B**, and Suppl. Table 1**). Hierarchical clustering based on enhancer acetylation readily separated B-cell cancers by both primary type and B-ALL subtype (Suppl. Fig. 1A). Enhancers active in B-ALL subtype-specific clusters were observed near previously identified subtype-specific signature genes^1^ (Figure 1C **and** Suppl. Fig 1C), suggesting that enhancers contribute to subtype-specific gene expression programs. *De novo* motif enrichment analysis of enhancer clusters that were hyperacetylated in specific B-ALL cell lines revealed strong enrichment for motifs of TFs known to have increased activity in the corresponding subtypes, including DUX4 (B-ALL with *DUX4* rearrangement), HLF (*TCF3-HLF*+ B-ALL), HoxA9 (B-ALL with *KMT2A* rearrangement), and STAT5 (*BCR-ABL1*+ and *BCR-ABL1*-like B-ALL; Figure 1D-E, Suppl. Fig 1D-E**, and Suppl Table 2**). Surprisingly, cluster 12 enhancers that were hyperacetylated in E-R+ B-ALL cell lines (n=4) and in the BCR-ABL1-like B-ALL cell line MUTZ-5 were highly enriched for a motif consisting of tandem repeats of the sequence “GGAA.” Comparing acetylation levels across GGAA repeats of varying lengths in the hg38 reference genome showed increased acetylation in these five cell lines for intervals containing at least three repeats, with stronger effects at intervals with greater than six tandem repeats (Figure 2A; Suppl Figure 2A). Similarly, analysis of ATAC-Seq data showed increased chromatin accessibility in the same five cell lines for intervals containing 6x GGAA repeats compared to other B-ALL cell lines (Figure 2B). Similar analysis performed on ATAC-Seq^14^ and H3K27ac ChIP-Seq^15^ data from primary B-ALL samples confirmed a strong association between GGAA repeats and an active enhancer-like chromatin state in E-R+ B-ALL (Figure 2C-D; Suppl. Figure 2B).

**Figure 1:**
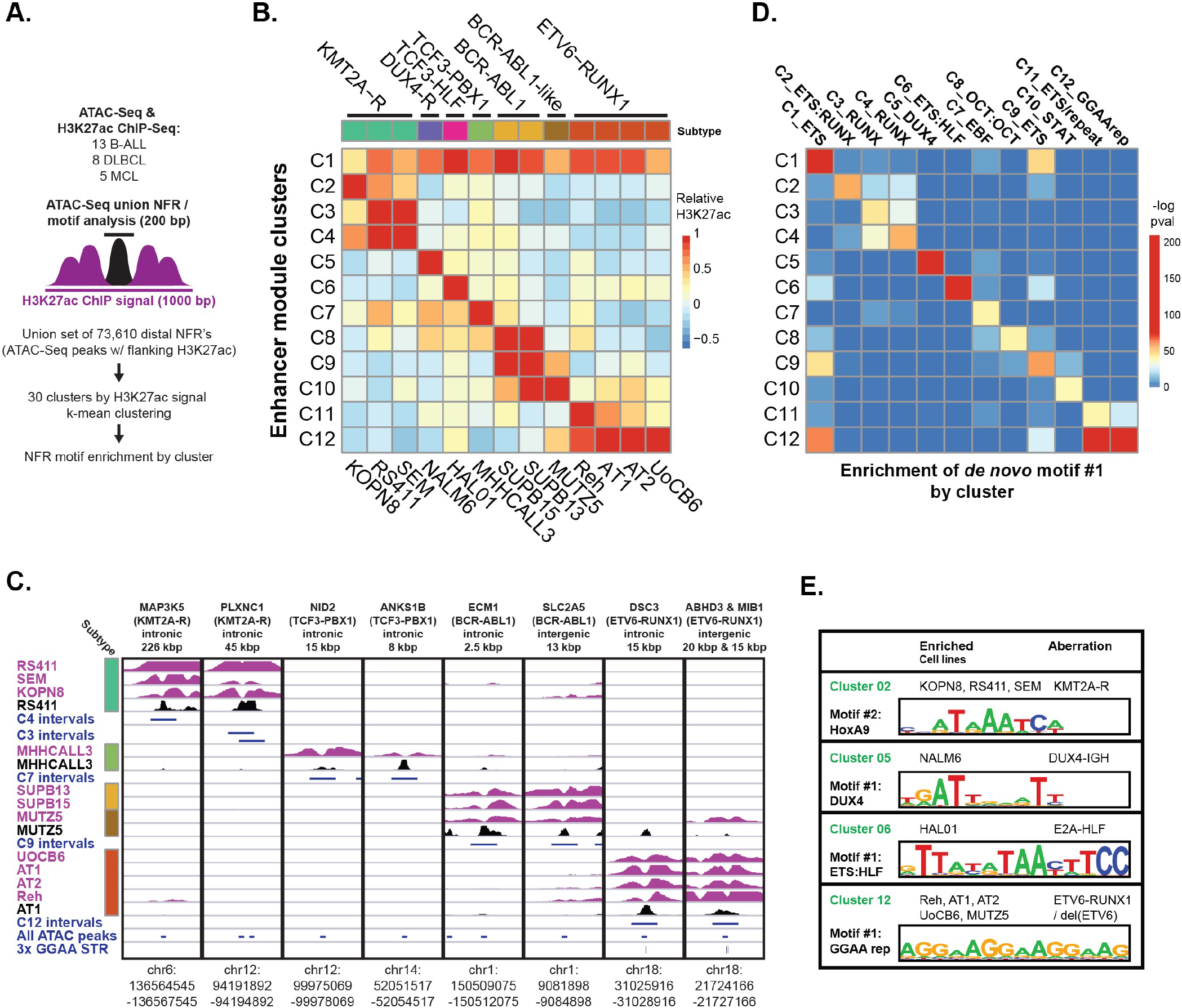
Identification of DNA motifs enriched in enhancers with subtype-specific activity. **A.** Strategy for identification of enhancer module acetylation clusters across 26 B cell cancer cell lines. **B.** Median acetylation signal in each B-ALL cell line for each of the 12 B-ALL-specific enhancer acetylation clusters (C1-C12), relative to all 26 cell lines (mature B-cell lines not shown). Genetic subtypes are listed at top and cell lines at bottom. **C.** H3K27ac ChIP-Seq and ATAC-Seq data for representative enhancers from subtype-specific clusters. Distance and position with respect to known B-ALL signature gene (Ross et al 2003)^1^ are listed at top, with the associated B-ALL subtype in parentheses. H3K27ac ChIP-Seq tracks are shown in purple (scale: 15 fragments per million, fpm) and ATAC-Seq tracks in black (scale 7.5 fpm). Intervals shown in blue correspond to the cluster-specific enhancer modules defined in (B) (1 kbp), union of all ATAC-Seq peaks (200 bp), and position of 3x GGAA tandem repeats. **D.** Significance of enrichment for the top *de novo* motif identified by HOMER in each of the clusters from (B), re-evaluated in all 12 clusters. **E.** Selected top *de novo* motifs identified in enhancer acetylation clusters.

**Figure 2:**
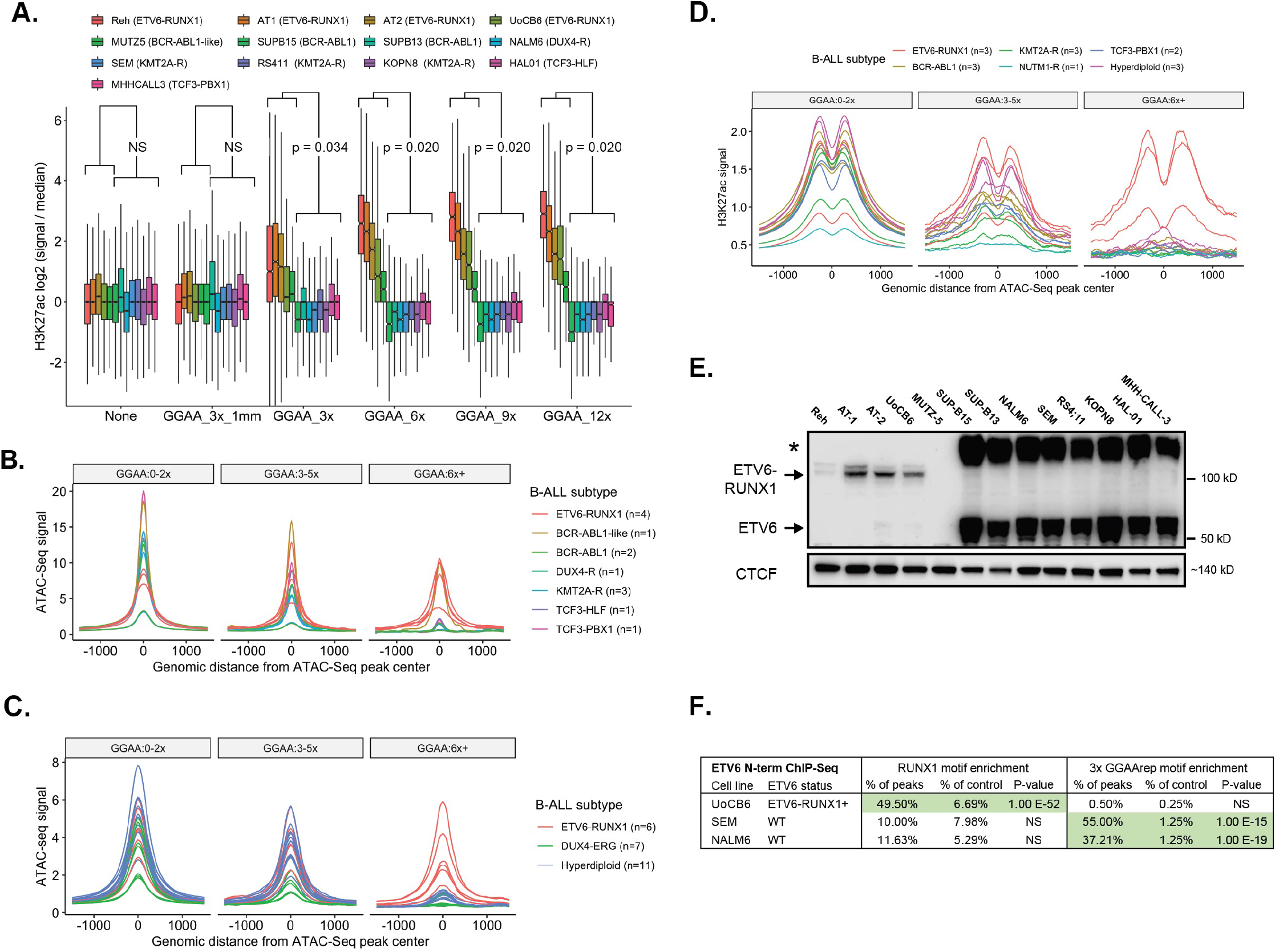
GGAA microsatellites show enhancer-like chromatin state in B-ALL with ETV6-RUNX1 and loss of wild-type ETV6. **A.** Boxplots showing relative H3K27ac levels flanking merged GGAA repeats and ATAC-seq peak-containing intervals identified across 13 B-ALL cell lines. Intervals were grouped according to the longest GGAA tandem repeat present. The “GGAA_3x_1mm” group contained a motif with a one-base mismatch to a 3x GGAA repeat. “None” indicates the set of ATAC-Seq peaks that do not contain a GGAA repeat by any criteria within 300 bp of the peak center. Results shown for Mann-Whitney U test of difference in means in ETV6-altered vs. ETV6-intact cell lines for each repeat class. **B.** Histogram of ATAC-Seq signal (reads per 10m total reads / bp / peak) from B-ALL cell lines of the indicated subtypes, centered on the union set of ATAC-seq peaks from those same samples. Peaks were grouped according to overlap with 3x or 6x GGAA tandem repeats. Note that the single BCR-ABL1-like cell line (MUTZ-5) is ETV6-null. **C.** Histogram of ATAC-Seq signal (reads per 10m total reads / bp / peak) from primary B-ALL samples of the indicated subtypes, centered on the union set of ATAC-seq peaks from those same samples. Peaks were grouped as in (B). **D.** Histogram of H3K27ac signal (reads per 10m total reads / bp / peak) from primary B-ALL samples of the indicated subtypes, centered on the union set of B-ALL cell line ATAC-seq peaks (n=13). Peaks were grouped as in (B). **E.** Immunoblot of nuclear extracts from 13 B-ALL cell lines with an antibody recognizing the N-terminal portion of ETV6 (Sigma, HPA000264). Arrows indicate bands at the expected molecular weight of ETV6 and E-R respectively. Asterisk indicates an apparent high-molecular weight form of ETV6. **F.** Enrichment for RUNX1 and 3xGGAA repeat motifs in ETV6 ChIP-Seq peaks (HOMER known motif analysis).

The E-R fusion transcription factor is thought to bind primarily to enhancers and promoters enriched for the RUNX1 motif (TGTGG)^16–18^, which does not resemble the enriched repeat sequence. Since the sequence GGA(A/T) forms the core of the binding motif for ETS family TFs^19, 20^, we hypothesized that aberrant acetylation of GGAA repeats in this subset of B-ALL might be related to deficiency of normal ETV6 repressor function. Western blot confirmed absence of wild-type ETV6 protein expression in nuclear extracts from MUTZ5 and all four E-R+ cell lines (Figure 2E). *ETV6* gene copy number analysis combined with *ETV6-RUNX1* single-fusion FISH studies indicated deletion of the non-rearranged copy of *ETV6* in all four E-R+ cell lines, and biallelic *ETV6* deletion in MUTZ5 (Suppl. Figure 2C **and Suppl. Table 1**). ChIP-Seq performed with an N-terminal ETV6 antibody yielded peaks that were significantly enriched for the RUNX1 motif, but not GGAA repeats, in the E-R+ B-ALL cell line UoCB6. In contrast, ETV6 ChIP-Seq performed in two ETV6 wild-type cell lines showed enrichment for GGAA repeats (albeit with relatively few genome-wide peaks), but not RUNX1 motifs (Figure 2F **and Suppl. Table 3**), supporting a model where GGAA repeats represent a direct binding target for wild-type ETV6, but not for the E-R fusion protein.

To further investigate ETV6 binding targets and functions, we used a doxycycline-inducible construct to express V5-tagged ETV6^WT^ or ETV6^R399C^, a variant associated with loss of DNA binding function and hematopoietic abnormalities^12^, in the E-R+ cell line Reh (Suppl. Fig. 3A). Expression of ETV6^WT^-V5, but not ETV6^R399C^-V5, significantly reduced growth of Reh cells compared to tagBFP transgene-expressing controls (Figure 3A). ChIP-Seq with a V5 antibody identified 2,343 significant binding peaks in ETV6^WT^-V5 expressing Reh cells, of which 80% overlapped genomic sites with at least 3 perfect GGAA tandem repeats (Figure 3B). To understand the effects of ETV6 on chromatin state, we performed replicate H3K27ac ChIP-Seq in Reh cells after doxycycline induction of ETV6^WT^, ETV6^R399C^, or tagBFP control. Expression of ETV6^WT^ was associated with deacetylation of histones flanking GGAA repeats (Figure 3C), with a stronger effect seen at sites with longer repeats and at sites where ETV6-WT-V5 binding was detected by ChIP-Seq. In contrast, expression of ETV6^R399C^ resulted in minimal acetylation changes at these same sites. These findings support the surprising conclusion that ETV6 chromatin repression activity is primarily directed at GGAA repeats upon restoration in ETV6-deficient B-ALL.

**Figure 3:**
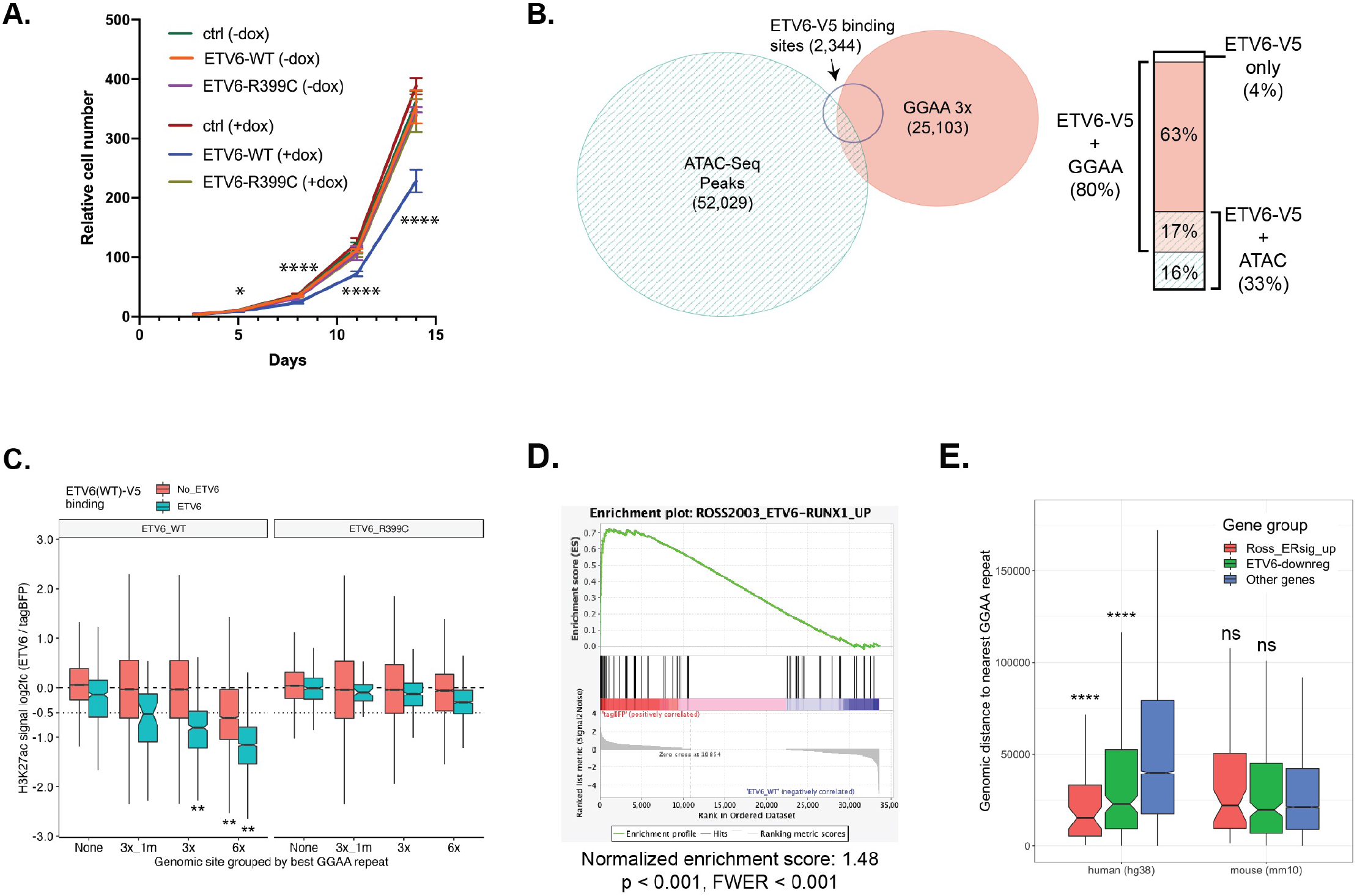
ETV6 inhibits leukemia cell growth, binds and deacetylates GGAA repeat enhancers, and activates E-R signature genes. **A.** Relative cell numbers for Reh cells stably transduced with DoxOn-ETV6-V5 constructs or DoxOn-tagBFP control and grown with or without 500 ng/ml doxycycline. Error bars show 95% CI of triplicate wells counted for each condition / time point. For each time point, log-transformed cell counts for ETV6-WT+dox were compared to each other sample by one-way ANOVA with Dunnett’s multiple comparison test. Comparisons at day 3 were non-significant and all comparisons were significant at later time points as follows. Day 5, padj < 0.05, Days 8, 11, and 14, padj < 10^-4^. **B.** Area-proportional Euler diagram showing overlap of genome-wide ATAC-Seq peaks in parental Reh cells, GGAA tandem repeats (at least 3x GGAA), and sites bound by ETV6-WT-V5 (V5 ChIP-Seq peaks) expressed in Reh cells. Bar plot at right details overlaps within the set of ETV6 binding target sites. **C.** Boxplots showing change in acetylation at ATAC-Seq peaks and / or GGAA repeats following induction of ETV6-WT or ETV6-R399C in Reh cells. Peaks were grouped by most stringent repeat class, and further divided by overlap with ETV6-WT-V5 ChIP-Seq peaks. Groups with log2 fold-change significantly less than −0.5 are indicated (Wilcoxon signed rank test with Holm-Bonferroni correction, ** padj < 10^-9^). **D.** Gene set enrichment analysis for upregulated E-R signature genes*^1^* among genes ranked by differential expression after induction of ETV6-WT-V5 or tagBFP (control) expression. **E.** Comparison of distance to the closest GGAA repeat (3x) for evolutionarily conserved human genes compared to their orthologs in mouse, separated into mutually exclusive sets of previously-defined E-R-upregulated signature genes (Ross 2003, red), non-ER signature genes that were downregulated by ETV6 expression in REH cells (green), and other genes (blue). Significance of gene sets vs “other genes” in each species calculated by Mann-Whitney U test with Holm-Bonferroni correction (**** = padj < 10^-7^)

We next investigated whether restoration of ETV6 would reduce expression of genes associated with GGAA repeat enhancers. Gene set enrichment analysis showed strong enrichment of known E-R+ B-ALL signature genes^1^ among ETV6-repressed genes (Figure 3D **and** Suppl. Figure 3B). E-R signature gene and other ETV6-repressed gene promotors were located significantly closer to the nearest GGAA repeat compared to non-regulated genes, but this relationship was lost for orthologs of those same genes in mice (Figure 3E). This finding is consistent with the poor overall conservation of GGAA repeat locations between humans and rodents (Suppl Fig 4), suggesting that the specific gene regulatory consequences of GGAA repeat enhancer formation in human cells are unlikely to be reproducible in mouse models.

To further evaluate whether ETV6 repression target genes identified in Reh cells were relevant in primary B-ALL, we focused on significantly repressed genes (padj < 0.001) associated with ETV6 binding sites within 200 kb from the promoter (**Suppl. Table 4)**. ETV6 binding sites showed a tendency for clustering near the promoters of ETV6-repressed genes (+/- 50 kb), while no such skew was observed for control genes (Figure 4A). For 78 ETV6-repressed genes, at least one associated ETV6 binding site showed decreased H3K27ac levels (p<0.05) upon ETV6-WT-V5 expression. We looked at expression of these genes in 118 primary B-ALL samples (11 E-R+ B-ALL and 107 other B-ALL) from the TARGET^21^ phase 2 cohort for which both RNA-Seq and genomic copy number segmentation data were available. Despite the relatively small size of this cohort, we found that 40 of the 78 ETV6 target genes were overexpressed in E-R+ samples compared to all other B-ALL (40 genes p < 0.05, 25 genes FDR-adjusted padj < 0.05, Suppl Fig 5). We combined the individually significant genes into a composite signature and compared E-R+ B-ALL with secondary ETV6 gene copy loss (n = 5) to E-R+ B-ALL without clear secondary ETV6 deletion (n = 6), demonstrating significantly higher expression of the ETV6-repressed gene signature (p = 0.015, t-test) in cases of E-R+ B-ALL with a second *ETV6*-inactivating event (Figure 4B-C). Of the 107 non-E-R+ B-ALL samples, four were strong outliers, with expression of ETV6-repressed genes comparable to that seen in E-R+ B-ALL. Three of the four outlier samples showed *ETV6* gene-inactivating events, a significant enrichment (1 of 93 *ETV6*-intact vs. 3 of 14 *ETV6*-abn, p = 0.0068, Fisher’s exact test).

**Figure 4:**
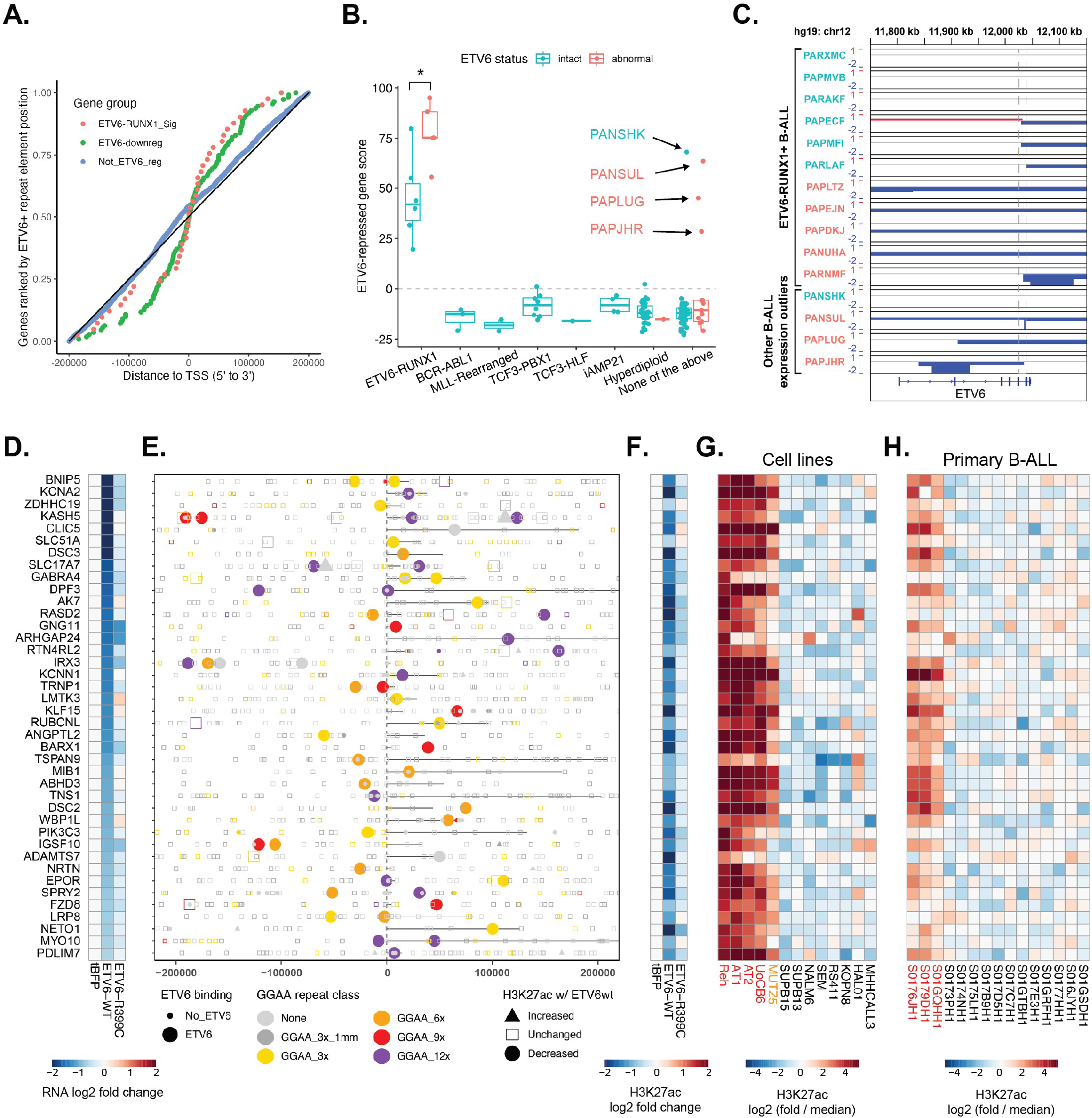
*ETV6*-repressed genes and enhancers are hyperactive in *ETV6*-altered B-ALL. **A.** Plot of distance from TSS to the nearest ETV6-WT-V5 binding site for genes repressed by ETV6 in REH cells from the set of E-R signature genes, other ETV6-repressed genes (“ETV6-downreg”), and a control set of expressed, non-ETV6-regulated genes. Only genes with an ETV6 binding site +/- 200 kbp from the promoter are included. **B.** Cumulative Z-score for expression of 40 ETV6-repressed gene with associated ETV6-repressed enhancers in primary B-ALL categorized by molecular subtype and ETV6 aberration status. Gene score was higher E-R+ B-ALL with secondary ETV6 aberrations (* p = 0.015, t-test). Four outlier samples with high expression of the ETV6-repressed gene score are indicated with black arrows. **C.** Segmentation tracks showing DNA copy number across the ETV6 locus in primary E-R+ B-ALL and in non-E-R+ B-ALL with outlier expression of ETV6-repressed genes (see B). Samples labelled in red were classified as “*ETV6* abnormal”, based on monoallelic deletions affecting *ETV6* 5’ to the chromosomal fusion hotspot in intron 5 (gray dashed lines), or deletions affecting *ETV6* 3’ to intron 5. **D-H.** Details of ETV6-repressed enhancer-gene pairs. **D.** Differential gene expression (RNA-Seq) for the indicated gene, normalized to tagBFP control. **E.** Position of analyzed intervals (union of ATAC-Seq, GGAA repeats, and ETV6-WT-V5 binding sites) within 200 kbp of the indicated gene TSS, coded by ETV6-WT-V5 binding status, best GGAA repeat class, and differential acetylation. Genes are oriented 5’ to 3’, with annotated gene bodies indicated by a black line. **F.** Differential acetylation of one element (associated with the gene listed in D) that shows ETV6 binding and significantly decreased acetylation in ETV6-WT-V5 expressing cells (H3K27ac log2 fold-change < −0.25, pval < 0.05), prioritized by repeat class (longest) and then distance to TSS (shortest, within 200 kbp). **G.** Relative acetylation of the element from F across 13 B-ALL cell lines (red = E-R+ / ETV6-null cell lines, orange = ETV6-null cell line). **H.** Relative acetylation of the element from F in 15 primary B-ALL samples (Blueprint project; red = E-R+ samples).

We examined the loci of ETV6-repressed genes validated in the TARGET cohort, and found that 38 of 40 were associated with an ETV6-bound GGAA repeat (at least 3xGGAA, Figure 4D-F, **Suppl. Table 4)**. One of the two exceptions, *CLIC5*, was associated with an ETV6-bound low-complexity element containing 23 individual GGAA sequences in 140 bp. Most of the associated ETV6-repressed repeat elements showed increased acetylation across the five ETV6-null B-ALL cell lines (Figure 4G), and many showed increased acetylation in primary E-R+ B-ALL compared to other subtypes (Figure 4H). Together, these findings indicate that overexpression of genes associated with ETV6-repressed microsatellite enhancers is a characteristic feature of E-R+ B-ALL, is accentuated in B-ALL with deletion of the second *ETV6* allele, and is seen in a subset of other B-ALL with *ETV6* inactivation events.

Next, we directly tested the functional relationship between specific ETV6-regulated GGAA repeats and their putative target genes. We designed sgRNAs to target six ETV6-regulated GGAA repeat enhancers via CRISPR-interference (CRISPRi) in Reh cells (Figure 5A). In each case, doxycycline-induced expression of a dCas9-KRAB repressor led to significant downregulation of the expected target gene, validating these genes as *bona fide* regulatory targets of GGAA microsatellite enhancers (Figure 5B). Functionally validated microsatellite enhancer activation targets include oncogenically significant genes such as *EPOR*, which encodes the erythropoietin receptor and confers increased STAT signaling and proliferation of primary E-R+ leukemia cells in the presence of erythropoietin^22, 23^, and *PIK3C3* (VPS34), a regulator of vesicle trafficking that has been reported to mediate dependence on autophagy in E-R+ B-ALL^24^.

**Figure 5:**
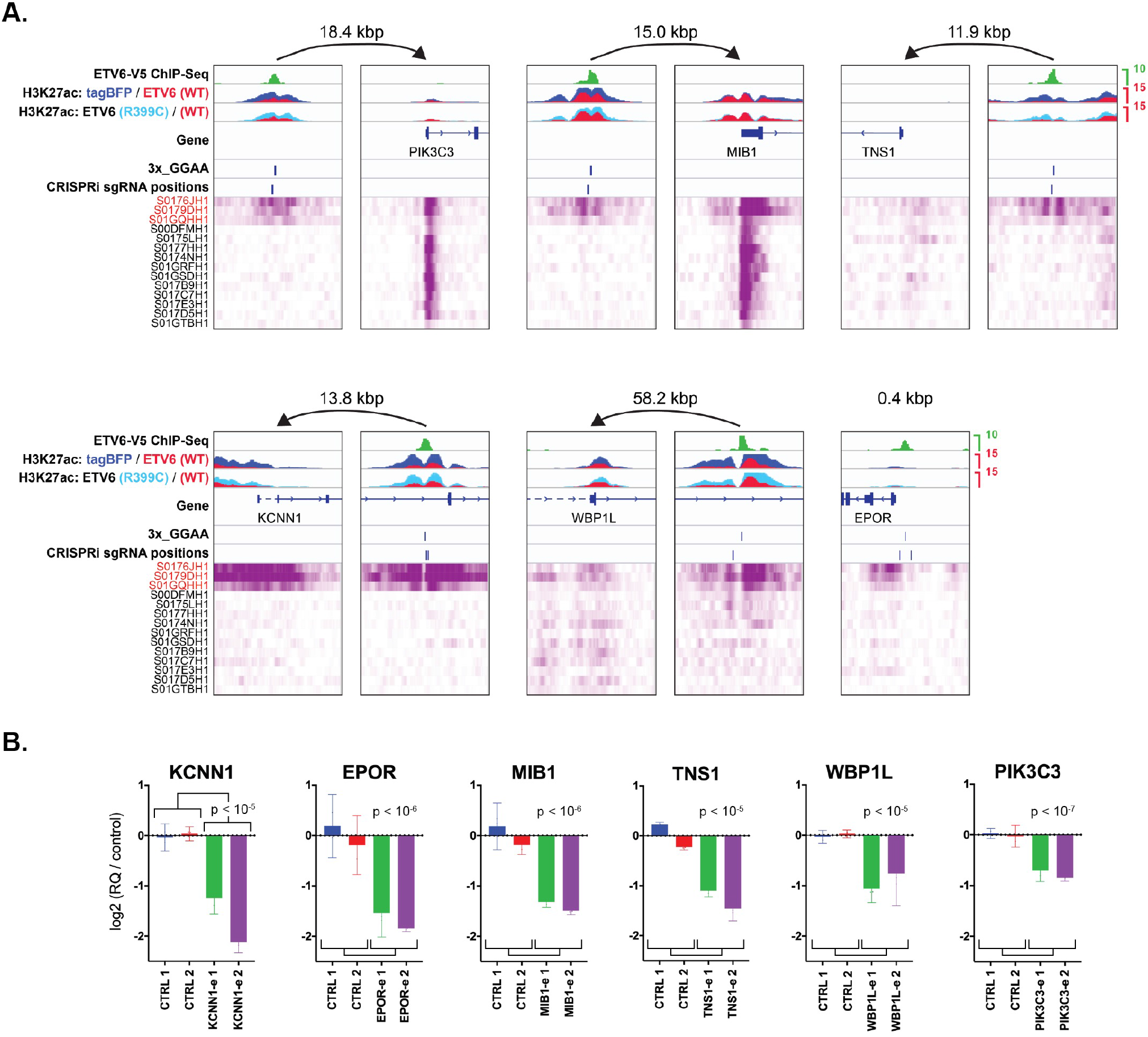
GGAA microsatellite enhancers are direct regulators of ETV6-RUNX1 signature gene expression. **A.** ChIP-Seq data from transgene-expressing Reh cells (top) and primary B-ALL samples (bottom) for selected E-R signature gene promoters and linked ETV6-regulated enhancers. ETV6-WT-V5 ChIP-Seq is from doxycycline-induced, transgene-expressing Reh cells. H3K27ac ChIP-Seq data was generated from tagBFP, ETV6-WT-V5, and ETV6-R399C-V5 cells, and overlays are color-coded as indicated. Also shown are positions of GGAA repeats called genome-wide in hg38 (3xGGAA), and sgRNAs used for CRISPRi in (B). Names of E-R+ primary B-ALL samples (Blueprint) are indicated in red. **B.** Relative transcript levels for genes shown in (A) 72 hrs after doxycycline induction of Reh cells expressing doxycycline-inducible dCas9-KRAB and transduced with control sgRNAs or sgRNAs targeting the indicated GGAA microsatellite enhancer. Gene expression is normalized to the average of the two control sgRNAs (error bars = 95% CI of PCR replicates). Significance calculated as t-test of combined replicates for both control sgRNAs versus both enhancer-targeting sgRNAs.

We next sought to identify TFs that contribute positively to GGAA repeat enhancer activation in *ETV6*-altered B-ALL. Only a minority of GGAA repeats in the hg38 reference genome are accessible and associated with substantial H3K27ac levels in E-R+ B-ALL cell lines (Suppl Fig 6A-B). Repeat-containing intervals with acetylation that was strong and specific to E-R+ B-ALL cell lines were significantly enriched in several classes of TF motifs (Figure 6A) compared to non-acetylated repeats, suggesting that binding of specific TFs in the vicinity of repeats might contribute to repeat enhancer activation. TFs that contribute to GGAA repeat enhancer activation likely overlap with those that regulate other B-cell enhancers, as these same motifs were also common in intervals that are strongly acetylated in all types of B-ALL, (Suppl Fig 6B-C). We were interested to note that the motif corresponding to the ETS activator ERG was particularly abundant near acetylated GGAA repeats. Both *ERG* and its homolog *FLI1*, are highly expressed in B-ALL, and the fusion oncoproteins EWSR1-ERG and EWSR1-FLI1 are known activators of GGAA microsatellite enhancers in the pediatric bone tumor Ewing sarcoma. However, DepMap CRISPR knockout screens show a substantial growth dependency on *ERG*, but not *FLI1*, for the E-R+ cell line Reh and most other B-ALL cell lines screened to date (Figure 6B **and** Suppl fig 6D).

**Figure 6:**
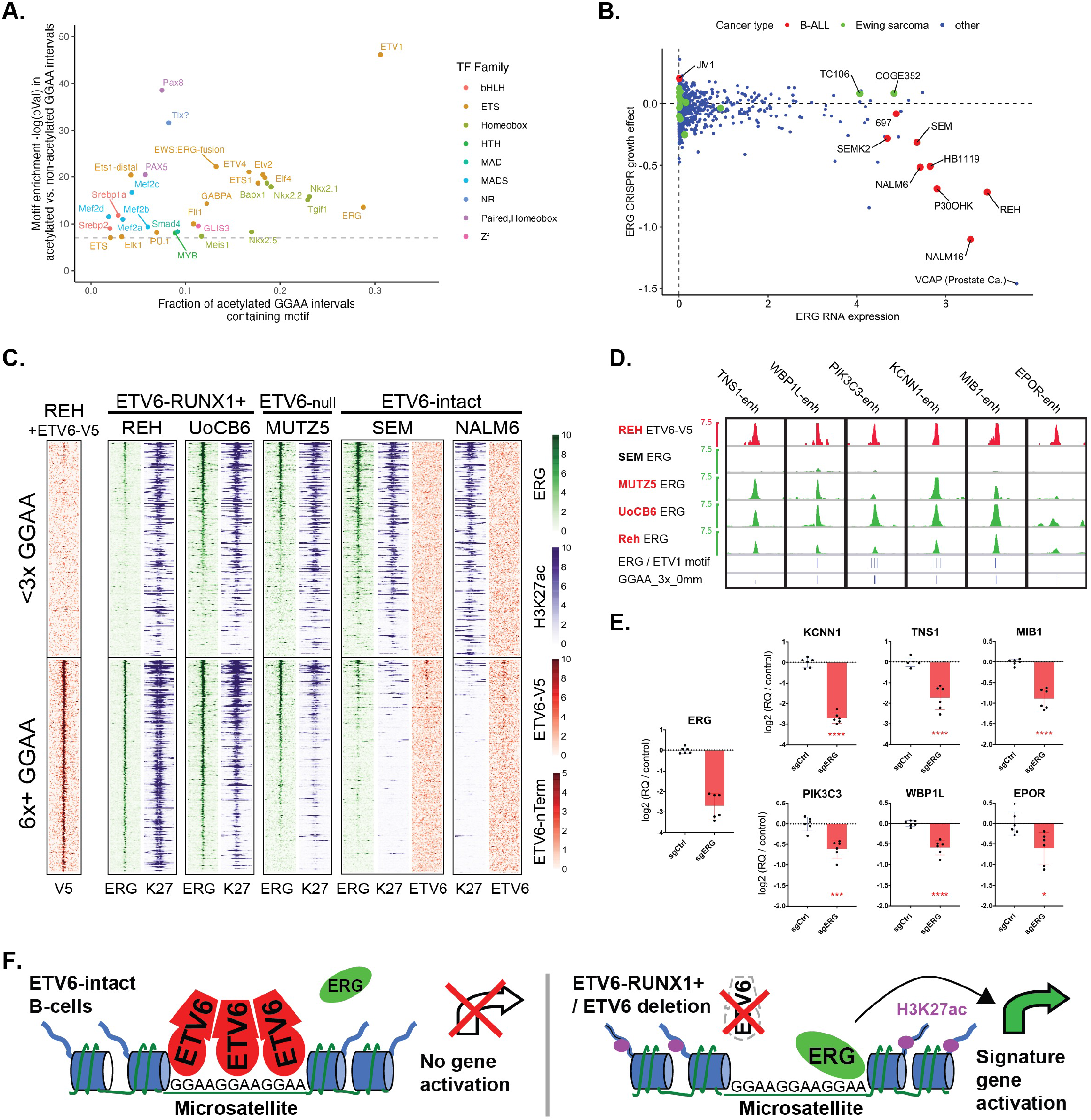
ERG contributes to GGAA repeat enhancer activity in ETV6-altered B-ALL. **A.** Enrichment of known TF motifs (HOMER motif library) in 200 bp intervals centered on GGAA repeats with selective strong acetylation in E-R+ B-ALL cell lines, compared to repeat intervals not associated with acetylation (see Supplementary figure 6B for thresholds used). All motifs with enrichment −log(pval) > 7 are shown. Note that motif analysis may have limited ability to discriminate between factors within a given family. **B.** DepMap data showing ERG expression and CRISPR knockout growth effect for B-ALL cell lines vs other cancer types. The *TMPRSS2–ERG+* prostate cancer cell line VCAP is also labeled. **C.** Heatmap of ERG, ETV6, and H3K27ac ChIP-Seq signal at repeat-containing (at least 6x GGAA) and non-repeat-containing (<3x GGAA) ATAC-Seq peaks in B-ALL cell lines. Peaks shown had H3K27ac fragment counts > 5 per million in at least one of 13 cell lines. The <3x GGAA group was randomly down-sampled to the same number of peaks as the 6x GGAA group. ATAC-Seq peaks were sorted according to the maximum ERG signal across the four ChIP-Seq datasets. V5 ChIP-Seq was performed in Reh cells induced to express ETV6-WT-V5; all other ChIP-Seq studies were performed in parental cell lines. **D.** ERG ChIP-Seq signal at CRISPRi-validated GGAA repeat enhancers shown in Figure 5. Positions of 3xGGAA repeats and predicted high-affinity motifs for ERG and ETV1 within +/- 250 bp of the central GGAA repeat (HOMER motifs and thresholds) are shown at bottom. **E.** Gene expression differences (qRT-PCR) 72 hours after doxycycline induction in Reh cells expressing doxycycline-inducible dCas9-KRAB and promoter-targeting sgRNA against *ERG* versus control (non-targeting) sgRNA. Values represent pooled PCR replicates from two separately conducted experiments (two-tailed t-test, * p < 0.05, *** p<0.001, **** p<0.0001). **F.** Model for role of ETV6 and ERG in regulation of GGAA repeat enhancers.

We performed ERG ChIP-Seq on B-ALL cell lines, which showed frequent ERG binding to acetylated GGAA repeats in the E-R+ B-ALL cell lines Reh and UoCB6, as well as the ETV6-null cell line MUTZ5, but far less binding to GGAA repeat enhancers in the ETV6-intact cell line SEM (Figure 6C). Interestingly, the small number of ERG-bound GGAA repeats that we did detect in SEM cells were often also bound by endogenous ETV6 repressor. We observed ERG binding to all six functionally validated GGAA repeat enhancers (Figure 6D), with four of the five enhancers showing predicted high-affinity ETS factor binding sites in addition to GGAA repeats. CRISPRi-mediated ERG knockdown resulted in substantially decreased transcript levels for all six genes, implicating ERG as a direct transcriptional activator of many repeat enhancer targets (Figure 6E-F). In contrast, we saw no consistent additional effect on the expression of GGAA repeat enhancer-regulated genes when we knocked down *FLI1*, either alone or in combination with *ERG* knockdown (Suppl. Figure 6E).

## Discussion

Distal regulatory elements play critical roles in oncogenic gene expression programs. While many well-characterized oncogene enhancers derive from evolutionarily conserved enhancers with normal functions in the cancer’s tissue of origin^25–27^, large-scale efforts to map cancer chromatin landscapes have revealed only partial overlap with enhancers known to be active in normal developing tissues^28^. The frequency with which human cancers rely on true *de novo* enhancers arising from non-conserved repeat elements remains an open question.

We found that the combination of ETV6 repressor insufficiency and ERG activator expression facilitate aberrant activation of GGAA microsatellite enhancers in a subset of B-ALL, and represent a key mechanism underlying the unique gene activation program of E-R+ B-ALL. Our findings suggest a mechanistic explanation for the occurrence of the E-R+ B-ALL gene expression program in rare cases of B-ALL that lack E-R but show *ETV6*-inactivating mutations and deletions^2, 11^. We saw some degree of GGAA repeat chromatin activation and repeat enhancer target gene overexpression in all primary E-R+ B-ALL datasets we examined, although not all E-R+ B-ALL cases show biallelic *ETV6* inactivation. *ETV6* haploinsufficiency, along with the known dominant-negative effects of E-R protein^29^ and other altered forms of ETV6 that lack the ETS domain^29, 30^ on wild-type ETV6 repressor function, may compromise silencing of repeats in cases without biallelic *ETV6* alteration. Further work is indicated to investigate a potential role for microsatellite enhancers in B-ALL occurring in the setting of germline *ETV6* mutations, which are reported to be heterogeneous in their biology and gene expression signatures,^13^ and in diverse (albeit rare) leukemias and solid tumors that bear rearrangements between *ETV6* and gene partners other than *RUNX1*^31, 32^, at least some of which involve biallelic *ETV6* inactivation^33–35^.

The *in vitro* DNA sequence affinity of the ETV6 DNA binding domain is similar to that of other endogenous human ETS factors, all of which recognize a core motif of GGA(A/T)^19, 20^. However, ETV6 (and its homolog ETV7) are distinct in their ability to oligomerize via their N-terminal PNT domain^36–38^, as the PNT domains present in ERG and many other ETS factors do not self-associate^20, 39^. The self-associating property of the ETV6 PNT domain has been shown to confer cooperative binding of ETV6 to DNA sequences containing two ETS binding sites *in vitro*^38^, and models of *ETV6* oligomers binding to greater numbers of low-affinity core binding sites have previously been proposed^37^. These properties may explain our unexpected observation that ETV6 shows a strong *in vivo* preference for binding to GGAA repeat elements rather than canonical ETS motif-containing enhancers in B-ALL.

Our findings imply that a significant function of ETV6 is to sustain or restore epigenetic silencing of GGAA repeats that can serve as low-affinity ETS binding sites, and could otherwise be prone to aberrant enhancer-like activity. While GGAA repeat enhancer activation is a well-documented function of ERG and FLI1 fusion oncoproteins in Ewing sarcoma^40, 41^, our findings indicate that the non-fusion form of ERG contributes to GGAA repeat enhancer activation in the absence of ETV6, and that repeats with adjacent high-affinity binding sites for ERG or other factors may be particularly prone to aberrant activation.

Identification of microsatellite enhancers as key drivers of the E-R+ B-ALL gene expression program has several other important implications. Genetic mouse models of E-R expression alone yield either absent or very rare B-ALL^42–45^, while B-ALL generated by the combination of E-R and transposon-mediated random gene disruption failed to recapitulate the transcriptional program of human E-R+ B-ALL and showed no selection for ETV6-inactivating second-hits^46^. Our findings suggest limitations of engineered mouse models and widely-used mouse pro-B cell lines such as Ba/F3 for studying the biology of E-R+ and *ETV6*-deficient B-ALL, since most GGAA microsatellites present in the human genome lack similarly-positioned orthologs in rodents. Notably, attempts to model Ewing sarcoma in mice have also yielded very limited results^40^. Because enhancer-like activity of GGAA microsatellites is not known to contribute to the biology of any normal human cell, further investigation of mechanisms required to sustain this phenomenon could yield appealing targets for selective therapeutic intervention.

## Methods

### Cell lines

See **Suppl. Table 1** for details of cell line source and validation. The identity of all B-ALL cell lines was verified by short tandem repeat (STR) profiling. All cell lines matched the expected profile from public databases (when available), and all cell line profiles were unique, with the exception of pairs of cell lines derived from the same donor (AT-1 and AT-2^47, 48^, and SUP-B13 and SUP-B15^49^), which showed mutually identical profiles as expected. All cell lines were cultured in RPMI 1640 + Glutamax supplemented with 10% fetal calf serum, penicillin/streptomycin, nonessential amino acids, 1 mM sodium pyruvate, and 55 μM 2-mercaptoethanol.

Cell line cytospins were evaluated using fluorescent in situ hybridization (FISH) probes for ETV6 (Chr12p13; SpectrumOrange; Abbott) and RUNX1 (Chr21q22; SpectrumGreen; Abbott) by standard techniques. Cell lines with one or more fusion signals per cell were considered positive for an E-R rearrangement, with 1F1R2G representing the expected pattern for a single balanced E-R rearrangement. Note that the ETV6 probe covers the 5’ portion of the ETV6 gene and a large region of chromosome 12 (486 kb). Absence of a red signal therefore indicates absence of a non-rearranged copy of ETV6, but a focal ETV6 deletion is not excluded by presence of a red signal.

Genomic copy number analyses of cell lines were carried out as previously described. Briefly, depth-coverages for the for the input lowpass WGS (ChIP-seq chromain input) libraries were quantified at a 50kb resolution (bins), the resulting data were segmented, and a probabilistic model^50, 51^ was fit to assign absolute copy number states to the observed coverages. The models were constrained assuming 100% neoplastic cellularity.

### ChIP-Seq

Protocols used for new ChIP-Seq datasets were similar to those previously described^52, 53^. Briefly, five million cells were cross-linked in PBS + 1% formaldehyde for 10 minutes at room temperature (histone marks) or 37°C (TFs), quenched with 1/20^th^ volume of 2.5M glycine, washed twice in cold PBS with protease inhibitors, and lysed in cold cytoplasmic lysis buffer (20 mM Tris-HCl ph8.0, 85 mM KCl, 0.5% NP 40 + PI). Nuclei were pelleted at 3000g, and resuspended in cold SDS lysis buffer (0.3% SDS for H3K27ac and 1% SDS for TFs, 10mM EDTA, 50mM Tris-HCl, pH 8.1 + PI) for 10 minutes. Nuclei were fragmented on a Q800R2 Sonifier (QSonica) as follows: three cycles of amplitude = 50, pulse times: 30s on / 30s off, total on time = 3:20 m, temperature = 8^0^C (histone marks) or two cycles of amplitude = 70, pulse times: 45s on / 15s off, total on time = 8:50 m, temperature = 4^0^C (TFs). Samples were diluted 1:3 in ChIP dilution buffer (0.01% SDS, 1.1% Triton X-100, 1.2mM EDTA, 16.7mM Tris-HCl, pH 8.1, 167mM NaCl +PI), and rotated at 4°C overnight with 2-5 ug of antibody (H3K27ac, Active Motif Cat #39133; V5 tag, ThermoFisher #R960-25; ETV6, Santa Cruz, # sc-166835x; ERG, Cell Signaling Technologies # 97249). Subsequent chromatin capture, washing, DNA elution, purification, and Illumina library preparation steps were performed as previously described. Libraries were sequenced on NextSeq High-output flow cell 75 cycles (2 x 38 bp paired end sequencing).

See Supplementary Table 1 for the list of new and previously published ChIP-Seq datasets used in this study, as well as reference to corresponding protocols used.^52, 53^ Datasets generated on a subset of cell lines with both new and old methods showed qualitatively equivalent results.

### ATAC-Seq

Nuclei were isolated from 50,000 cells for each sample using Nuclei EZ prep-Nuclei Isolation Kit (Sigma-Aldrich, USA). The transposition reaction mix (25μl of 2X TD buffer, 2.5μl of Tn5 transposase (Illumina, San Diego, CA, USA), 15ul of PBS and 7.5μl of nuclease free water) was added to nuclei and incubated at 37°C for 1hour in an orbital shaker at 300RPM. 50uL Qiagen buffer PB was added to each sample to stop the reaction and DNA was isolated with AMPure XP beads (Beckman Coulter). 15 cycles of PCR were performed with transposed DNA using the dual index primers and NEBNext PCR Master Mix, followed by AMPure XP purification. After quantification and fragment size analysis, libraries were sequenced on Illumina Nextseq with 2 x 38 bp paired-end sequencing.

### ETV6 transgene experiments

The lentiviral vector DoxON-ETV6-V5-GFP was a kind gift from the lab of Dr. Arul Chinniayan. The ETV6 coding sequence was cloned into pCW57.1 (Addgene cat no. 41393) modified as previously described to incorporate a GFP reporter^54^. DoxON-tagBFP was generated from that vector by restriction cloning tagBFP in place of ETV6 after digestion with BstBI and BmtI. DoxON-ETV6(R399C)-V5-GFP was generated with the Q5 site-directed mutagenesis kit (NEB) per manufacturer’s protocol.

Lentivirus was produced in 293T cells by standard protocols. To generate uniform doxycycline-inducible populations, Reh cells were transduced with DoxON constructs via spinfection at 2250 rpm for 90 minutes at 37°C in the presence of 6 µg/ml polybrene and sorted for GFP+ cells on a BD MoFlo Astrios EQ. Uniformly sorted Reh cell populations were induced with 500 ng/ml doxycycline (dox) for 48 hours prior to harvest for western blot or V5 ChIP-Seq. Induction was performed in duplicate for H3K27 ChIP-Seq or triplicate for RNA-Seq. To determine the effect of transgene expression on cell growth, each population was plated at equal density in triplicate wells with and without doxycycline. Cells were counted every 4 days on a DeNovix CellDrop BF counter with trypan blue staining, and re-plated at equal densities.

### RNA-Seq

RNA was isolated via RNEasy columns with on-column DNAase digestion. RNA-seq libraries were generated with the NEBNext Ultra II Directional RNA Library Prep Kit for Illumina per manufacturer’s instructions and sequenced on an Illumina Nextseq with 2 x 38 bp paired-end sequencing.

### Western Blotting

Western blotting of nuclear extracts or whole cell extracts was performed by standard methods using antibodies specific for the N-terminal portion of ETV6 (Sigma, # HPA000264; Santa Cruz, #sc-166835x), CTCF (Cell Signaling, #3418), and actin (Santa Cruz, #sc-8432).

### Chromatin data analysis

Paired-end ChIP-Seq and ATAC-Seq reads were aligned to hg38 using BWA-ALN (v 0.7.17) and filtered to remove PCR duplicates and read-pairs mapping to >2 sites genome-wide. Display files were generated with deeptools bamCoverage and visualized with IGV. Scaling for all ChIP-Seq and ATAC-Seq tracks in figures is equal to local paired-end fragment coverage x (1,000,000/ totalCount). ERG, endogenous ETV6, and ETV6-V5 ChIP-Seq peak calling was performed with HOMER findPeaks using the ‘factor’ style and FDR < 0.001 (for ERG), FDR < 0.01 for ETV6-V5, and FDR <0.05 for endogenous ETV6. ATAC-Seq peak summits were identified with MACS2 using default parameters. All peak sets were post-filtered against hg38 blacklist regions (available at https://github.com/Boyle-Lab/Blacklist/blob/master/lists/hg38-blacklist.v2.bed.gz).

To analyze H3K27ac ChIP-Seq signal associated with individual enhancer modules, we resized ATAC-Seq peaks to 200 bp around MACS2 peak summits, discarded peaks with low signalValue (<5), and then used GenomicRanges ‘reduce’ and ‘resize’ to generate consensus union ATAC-Seq peak sets of 200 bp intervals for the samples of interest (26 B-cell cancer cell lines or 13 B-ALL cell lines dependening in the analysis). HOMER annotatePeaks was used to annotate union ATAC-Seq peaks with normalized H3K27ac ChIP-Seq signal from the relevant cell lines in a 1000 bp window around each interval center.

To identify clusters of enhancers with correlated acetylation levels across the 26 B cell cancer cell lines, we filtered out ATAC-Seq union peak intervals located < 2kb upstream or < 1kb downstream of an annotated TSS, as well as intervals associated with low H3K27ac signal in all cell lines. Acetylation signal was square root-transformed, centered, and scaled by genomic region across all cell lines. K-means clustering (k=30) was used to identify enhancer clusters. HOMER findMotifsGenome was used to identify both known and *de novo* enriched TF motifs in each cluster, with the set of all enhancers used as a background (option -b). The top two *de novo* motifs identified in each B-ALL-specific cluster were then used as custom known motifs to calculate enrichment in all B-ALL-specific clusters. For known motif enrichment analysis of endogenous (N-terminal) ETV6 ChIP-Seq peaks, the custom “GGAA_3x_0mm” motif described below was appended to the Homer known motif library.

### Identification of chromatin features and genes associated with GGAA repeat intervals

We used HOMER seq2profile.pl to generate HOMER custom motif files corresponding to a specified number of GGAA tandem repeats and permitted number of mismatches, e.g. motif “GGAA_3x_1mm” corresponds to a genomic sequence with no more than one mismatch to the sequence “GGAAGGAAGGAA”, while “GGAA_3x” corresponds to exact matches to that same genomic sequence. We then used HOMER scanMotifsGenomeWide.pl to identify all occurrences in the hg38 reference genome for the motifs GGAA_3x_1mm, GGAA_3x, GGAA_6x, GGAA_9x, and GGAA_12x. We used HOMER mergePeaks to merge identified motifs into uniform 200 bp genomic intervals (-d 200), each of which was centered on one or more motif instances and was annotated with the most stringent contained motif. This set of annotated repeat-containing intervals was then overlapped with the union set of ATAC-Seq peaks in 13 B-ALL cell lines. HOMER annotatePeaks was then used to annotate each interval in the union ATAC-Seq / GGAA repeat set with normalized H3K27ac ChIP-Seq signal in an 800 bp window.

To generate histograms of ATAC-Seq or H3K27ac ChIP-Seq signal associated with non-repeat and repeat-containing nucleosome-free regions, we assigned each peak in the union set of distal ATAC-Seq peaks from 13 B-ALL cell lines or 24 primary B-ALL samples (Diedrich et al 2021, re-processed as described above) to one of three groups based on whether it contained a 6x GGAA motif, 3x (but not 6x) GGAA motif, or neither. HOMER annotatePeaks (-hist 25 -size 3000) was used to generate normalized histograms for the appropriate ATAC-Seq or H3K27ac ChIP-Seq datasets for each of the three repeat groups.

To analyze chromatin effects of ETV6 restoration in Reh cells, Reh ATAC-Seq peaks were merged into a union interval set with ETV6-WT-V5 peaks and GGAA repeat-containing intervals (defined as above). Normalized H3K27ac ChIP-Seq signal from Reh cells expressing tagBFP, ETV6-WT-V5, and ETV6-R399C-V5 (two replicates each) were calculated in an 800 bp window around each peak. DESeq2 was used to calculate log2FoldChange values for acetylation associated with each peak, using default parameters with apeglm shrinkage. ggplot2 was used to display boxplots for differential acetylation data according to the repeat and ETV6 binding status of each interval.

RNA-Seq transcript levels for DoxON tagBFP, ETV6-WT-V5, and ETV6-R399C-V5 expressing cells were quantified with Salmon and collapsed to gene level (Ensembl, Feb2014) with tximport. Differential gene expression analysis was performed with DESeq2. The ROSS2003_ETV6-RUNX1_UP gene set was derived from Ross et al 2003^1^, Supplemental Information, section II “Top 100 chi-square probe sets selected for TEL-AML1, decision tree format”, including all genes with HD>50 above mean that could be successfully converted to Ensembl 2014 gene symbols. For gene set enrichment analysis, normalized gene-level RNA-Seq counts for tagBFP and ETV6-WT triplicates were exported by DESeq2 and converted to .gct format. GSEA_4.1 software was then used to calculate enrichment for the ROSS2003_ETV6-RUNX1_UP gene set.

To link differentially expressed genes to candidate regulatory elements, runSeq2gene (Bioconductor package: seq2pathway) was used to link each interval in the Reh ATAC-Seq / ETV6-V5 ChIP-Seq / GGAA repeat union interval set to each hg38 Ensembl 2014 gene TSS within 200 kbp. Intervals were annotated (HOMER annotatePeaks) with normalized acetylation signal for ETV6 transgene or tagBFP control transgene-expressing cells, 13 B-ALL cell lines, and Blueprint primary B-ALL samples.

For genome-wide comparison of GGAA repeat element acetylation and motif associations in ETV6-RUNX1+ B-ALL, ETV6-intact B-ALL, and Ewing sarcoma, HOMER annotatePeaks was used to annotate hg38 genome-wide repeat-containing intervals (as defined above) with normalized H3K27ac ChIP-Seq signal (800 bp window) from B-ALL cell lines plus the Ewing sarcoma cell lines A673 and SKNMC (data from Riggi et al 2014^41^). B-ALL cell lines were grouped as ETV6-null (Reh, AT-1, UoCB6, and MUTZ-5; AT-2 was omitted due to shared origin with AT-1), or ETV6-intact (SUP-B15, NALM-6, SEM, RS4;11, KOPN-8, HAL-01, and MHH-CALL-3; SUP-B13 was omitted due to shared origin with SUP-B15), and median acetylation values determined for each interval in each group. Intervals with median acetylation value log2(tags per million+1) < 2 for both the ETV6-null and ETV6-intact groups were defined as being “non-acetylated”, intervals with median acetylation value log2(tags per million+1) > 4 for both the ETV6-null and ETV6-intact groups were defined as having “shared acetylation” and intervals with log2(tags per million+1) < 4 in ETV6-intact, >4 in ETV6-null, and log2(ETV6-null tags per million+1) −log2(ETV6-intact tags per million+1) > 2 were defined as “ETV6-null-specific acetylation”. Homer findMotifsGenome was then used to determine enrichment of known TF motifs in the ETV6-null-specific acetylation intervals or shared acetylation intervals (200 bp window), versus the non-acetylated intervals used as a custom background (option -b).

To generate heatmaps of ETV6, ERG, and H3K27ac ChIP-Seq signal, the union set of ATAC-Seq peaks from 13 B-ALL cell lines was annotated with H3K27ac signal for 13 B-ALL cell lines (800 bp window) and filtered to retain peaks with normalized H3K27ac signal >5 fragments per million in at least one cell line. Peaks were then annotated with signal profiles from each ChIP-Seq dataset using HOMER annotatePeaks with options -size 8000 -hist 20 -ghist. Groups of peaks containing 6x GGAA repeats or no repeats (<3x GGAA) were retained, with the latter group randomly downsampled such that each group had equal numbers of peaks. Peaks were then sorted according to maximum ERG ChIP-Seq signal (400 bp window) across four cell lines for which ERG ChIP-Seq was performed.

### Primary B-ALL gene expression analysis

Uniformly processed TARGET Phase II B-ALL data was downloaded from cBioPortal, including gene expression Z-scores, hg19 genomic copy number abnormality segmentation, and sample metadata including *ETV6-RUNX1* FISH results and molecular subtype. Samples from unique patients with both RNA-seq and CNA segmentation data available (122 total) were analyzed. Molecular subtypes were used as provided except that the groups “Trisomy of both chromosomes 4 and 10” “Hyperdiploidy without trisomy of both chromosomes 4 and 10” and “Hyperdiploid; status of 4 and 10 unknown” were merged into a single “Hyperdiploid” group. Samples were assigned an “abnormal” ETV6 status if segmentation data showed monoallelic or biallelic deletion of any portion of the ETV6 genes, except for three E-R+ samples for which monoallelic deletion of *ETV6* on the 3’ side of the 5^th^ intron could represent loss of the der(12)t(12;21) chromosome without affecting the intact ETV6 gene. As these three ambiguous samples showed repeat enhancer gene signature scores intermediate between the samples with no ETV6 deletions and those with definitive secondary ETV6 deletions, including them in either the ETV6-intact or ETV6-deleted groups did not affect the statistical significance of our conclusions.

To identify a signature of ETV6-repressed genes, we identified genes that met the following criteria in our Reh ETV6-WT-V5 overexpression experiments: RNA log2 fold-change (tagBFP / ETV6-WT-V5) < −0.5, padj <0.001, and RNA log2 fold-change (tagBFP / ETV6-WT-V5) < log2 fold-change (tagBFP / ETV6-R399C-V5). We further filtered for genes linked to an ETV6 binding site within 200 kbp of the promoter that showed decreased H3K27ac signal in ETV6-WT-V5 vs tagBFP (1kb window, H3K27ac log2 fold-change < −0.25, pval < 0.05). 72 genes met these criteria, of which 40 were overexpressed (one-tailed t-test of RNA-Seq Z-scores, p < 0.05) in E-R+ B-ALL versus all other B-ALL in TARGET Phase II data.

### Comparative genomic analysis

The following approach was used to compare the relationship between GGAA repeats and gene promoters across mammalian species. HOMER scanMotifsGenomeWide was used to independently identify GGAA repeat-containing intervals in the hg38 (human), panTro5 (chimpanzee), Mmul10 (rhesus macaque), and mm10 (house mouse) genomes. UROPA^55^ was used to annotate the distance from each repeat to the start sites of all genes within 1 Mbp, using .gtf gene annotation files from ENSEMBL version 102. Gene-repeat linkages were filtered to retain only ENSEMBL genes with annotated homologs across all 4 species in the ENSEMBL 102 Biomart database. For pairwise comparisons between species, further filtering retained only the pair of gene homolog-repeat linkages with the shortest genomic distance in human, and only one pair of gene homologs per HUGO gene symbol, selected for the least difference in genomic distance from gene homolog to repeat between the two species.

### CRISPR-interference

To design sgRNAs targeting GGAA microsatellite enhancers, we used FlashFry^56^ to identify and score all candidate sgRNAs in a 2kb window around repeats of interest. Candidates were kept that met the following scoring criteria: Doench2014OnTarget > 0.1, Hsu2013 > 50, JostCRISPRi_specificityscore > 0.1, dangerous_GC == “NONE”, dangerous_polyT == “NONE”, dangerous_in_genome == “IN_GENOME=1”, otCount < 500. The final sgRNAs used for experiments were selected on the bases of shortest distance to GGAA repeat and highest Doech2014 on-target score (see Supplementary Table 1). Complementary oligonucleotides encoding sgRNA sequences plus appropriate overhangs were annealed and cloned into BsmBI-digested sgOpti (Addgene #85681).

A CRISPRi-ready Reh cell population with dox-inducible dCas9-KRAB and a GFP reporter (Reh-CiG) was generated as follows. Reh cells were transduced with lentivirus produced from TRE3-KRAB-dCas9-IRES-GFP and pLVX-EF1alpha-Tet3G vectors. Cells were serially sorted for GFP+ cells after doxycycline induction, for GFP-negative cells without doxycycline induction, and again for GFP+ cells after doxycycline induction.

For enhancer-targeting sgRNA experiments, Reh-CiG cells were transduced with control and repeat enhancer-targeting sgRNA lentivirus by spinfection. Cells were treated 48 hours after transduction with 1 ng/ml puromycin and 100 ng/ml doxycycline, and were harvested 5 days after transduction for RNA extraction and qRT-PCR.

Optimized promoter-targeting sgRNA sequences for knockdown of *ERG* and *FLI1* were selected from the “Dolcetto” genome-wide human CRISPRi library^57^. For knockdown of ERG and / or FLI1, variants of the sgOpti vector were generated by cloning tagBFP (sgMW-tagBFP) or tagRFP (sgMW-tagRFP) into BamHI and MluI-digested sgOpti in place of the PuroR gene. An ERG-targeting sgRNA sequence or non-targeting control was cloned into sgMW-tagRFP and a FLI1-targeting sequence was cloned into sgMW-tagBFP. Reh-CiG cells were transduced with appropriate combinations of control, ERG, FLI1, or ERG+FLI1 targeting sgRNA lentivirus and were flow sorted to ensure uniform expression of the appropriate fluorescent reporter(s). Cells were then induced with 500 ng/ml doxycycline for three days prior to RNA harvest and qPCR analysis.

## Supporting information

Suppl table 1 - Sequences and reagent validation

Suppl table 2 - HOMER motif analysis of clusters

Suppl table 3 - Summary of TF ChIP-Seq results

Suppl table 4 - ETV6-regulated gene-enhancer linkages

## Data availability

New sequencing datasets produced for this work are available at GEO under accession number GSE186942. Previously published data is available under accession numbers GSE69558 and GSE97541.

## Acknowledgements

We thank M. Le Beau and E. Davis for validating and providing B-ALL cell lines, D. Boyer for assistance with FISH, and A. Chinniayan, X. Wang, J. Kidd, P. Hsu, and E. Lawlor for helpful discussions. R.J.H.R acknowledges support from the National Cancer Institute (K08-CA208013) and a Hollis Brownstein Research Grant from the Leukemia Research Foundation. J.W.G acknowledges support from NIH training grant T32HL007622. A.C.M. acknowledges support from NIH training grant T32CA140044. The results published here are in whole or part based upon data generated by the Therapeutically Applicable Research to Generate Effective Treatments (https://ocg.cancer.gov/programs/target) initiative, phs000218. The data used for this analysis are available at https://portal.gdc.cancer.gov/projects.

## Supplementary figure legends

**Supplementary Figure 1:**
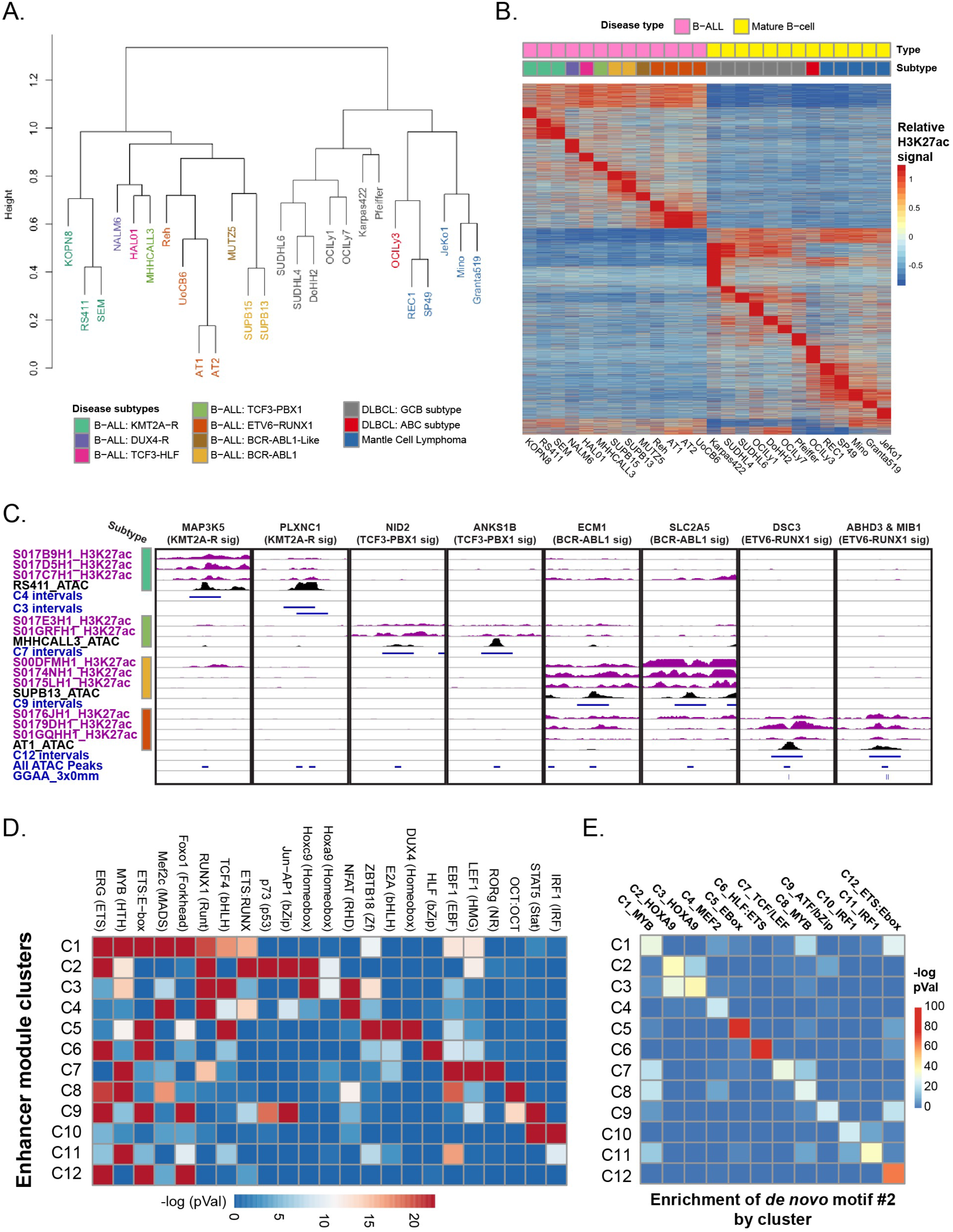
Additional details of B cell cancer cell line enhancer module clustering and motif analysis. **A.** Hierarchical clustering of 26 B-cell cancer cell lines based on Pearson distance of H3K27ac signal associated with 73,610 distal ATAC-Seq peaks. **B.** Relative H3K27ac ChIP-Seq signal flanking a union set of 73,610 distal ATAC-seq peaks across 26 B cell cancer cell lines, clustered by k-means clustering (k=30). **C.** Genomic regions showing the same B-ALL subtype-specific enhancers shown in Figure 1C, but with H3K27ac ChIP-Seq data from primary B-ALL samples (Blueprint Consortium) of the indicated subtypes. Intervals and B-ALL cell line ATAC-Seq are as described in Figure 1C. Subtypes are color-coded as in Figure 1B and Supplementary Figure 1A-B. **D.** Known motif enrichment analysis (HOMER library, selected motifs shown) for the 12 B-ALL-specific enhancer clusters. **E.** Significance of enrichment for the second most enriched *de novo* motif identified by HOMER in the 12 B-ALL-specific enhancer clusters.

**Supplementary Figure 2:**
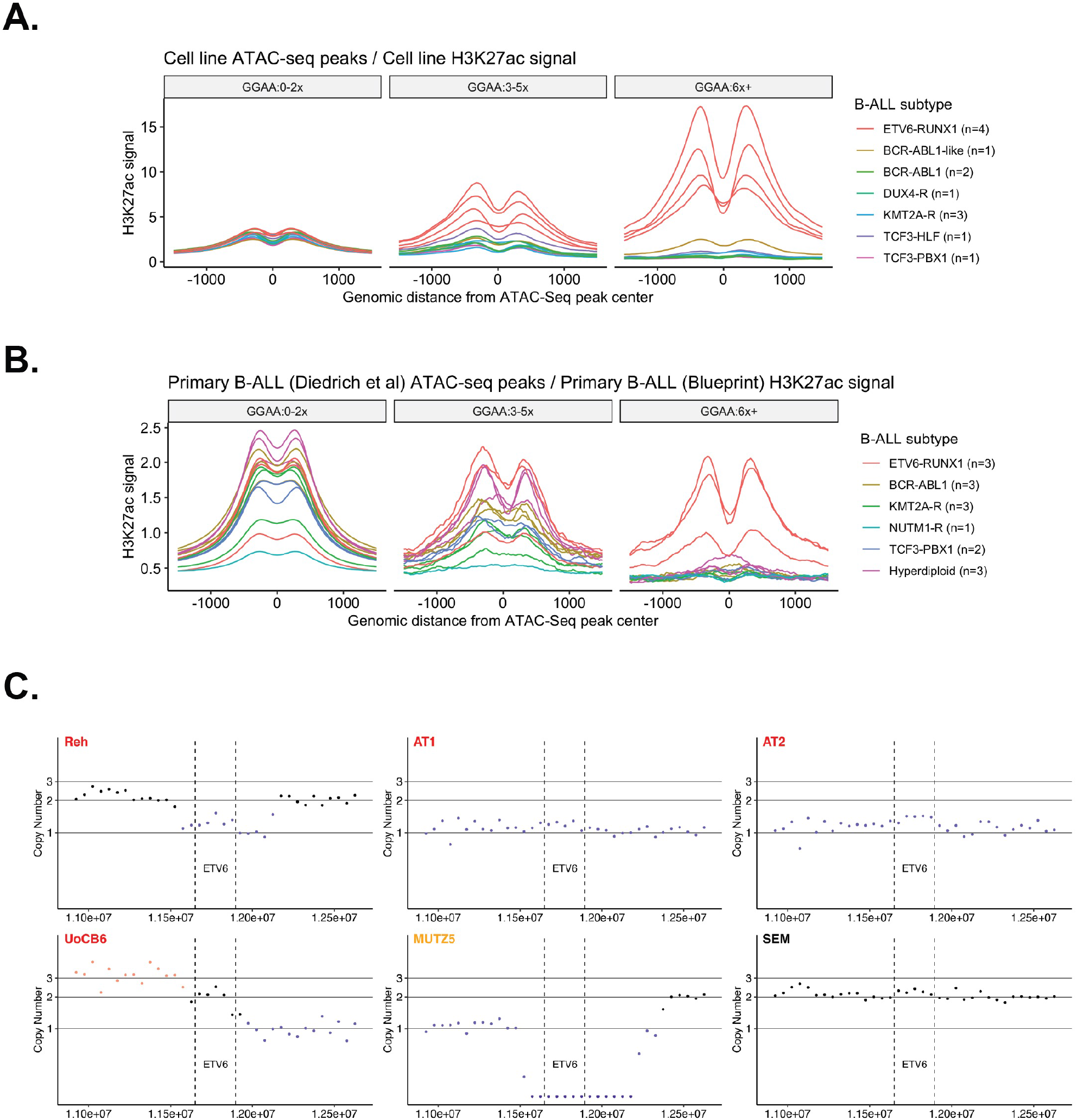
Additional analyses showing increased acetylation associated with GGAA repeats in ETV6-RUNX1+ and ETV6-null B-ALL. **A.** Histogram of H3K27ac signal (reads per 10m total reads / bp / peak) from B-ALL cell lines of the indicated subtypes, centered on the union set of B-ALL cell line ATAC-seq peaks (n=13). Peaks were grouped by overlap with repeat elements as in Figure 2B-D. Note that the single BCR-ABL1-like cell line (MUTZ-5) is ETV6-null. **B.** Histogram of H3K27ac signal (reads per 10m total reads / bp / peak) from primary B-ALL samples of the indicated subtypes, centered on the union set of ATAC-seq peaks from a different set of primary B-ALL samples (n=24). Peaks were grouped by overlap with repeat elements as in Figure 2B-D. **C.** DNA copy number abnormalities imputed across the ETV6 locus from ChIP-Seq input chromatin in E-R+ cell lines (red titles) and MUTZ5. ETV6 gene boundaries shown in dotted lines. All other cell lines were imputed as 2n across this locus (SEM shown as representative).

**Supplementary Figure 3:**
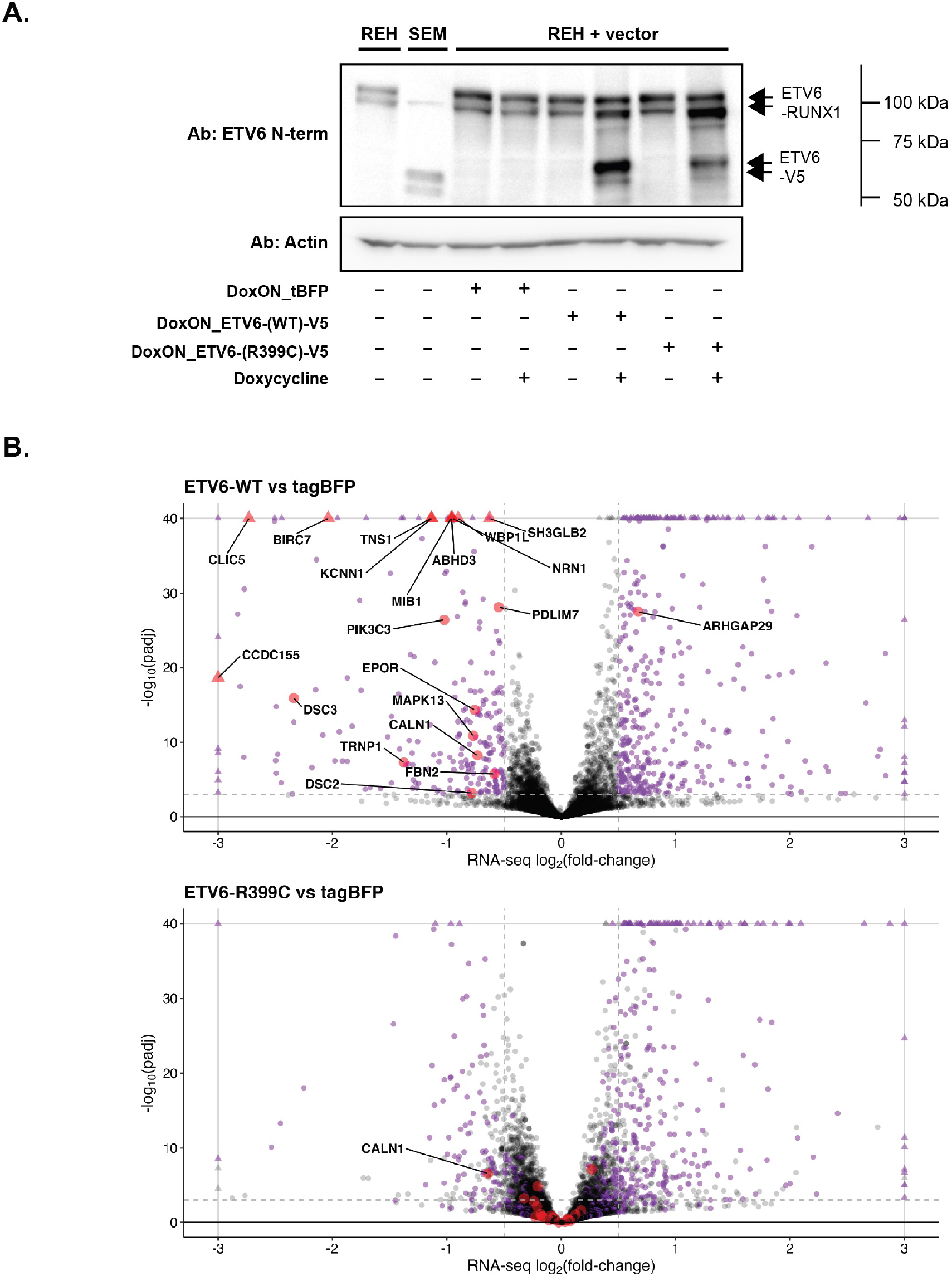
Additional details of ETV6 re-expression in the Reh cell line. **A.** ETV6 expression (n-terminal antibody) in parental B-ALL cell lines and Reh cell lines transduced with doxycycline-inducible ETV6-V5 constructs. **B.** Volcano plots showing differential gene expression for Reh cells overexpressing ETV6-WT-V5 or ETV6-R399C-V5 relative to tagBFP. Data points with x or y values outside of the plotted area are shown as triangles. All *E-R*-up signature genes as defined in Ross 2003 that showed significant differential expression in ETV6-WT-V5-expressing cells (log2fold change < −0.5 or > 0.5, FDR-adjusted padj < 10^-3^) are labeled and marked with large red dots. All other differentially expressed genes are colored purple. Genes in the lower plot (ETV6-R399C-V5 vs tagBFP) are colored as defined by the ETV6-WT-V5 vs tagBFP comparison.

**Supplementary Figure 4:**
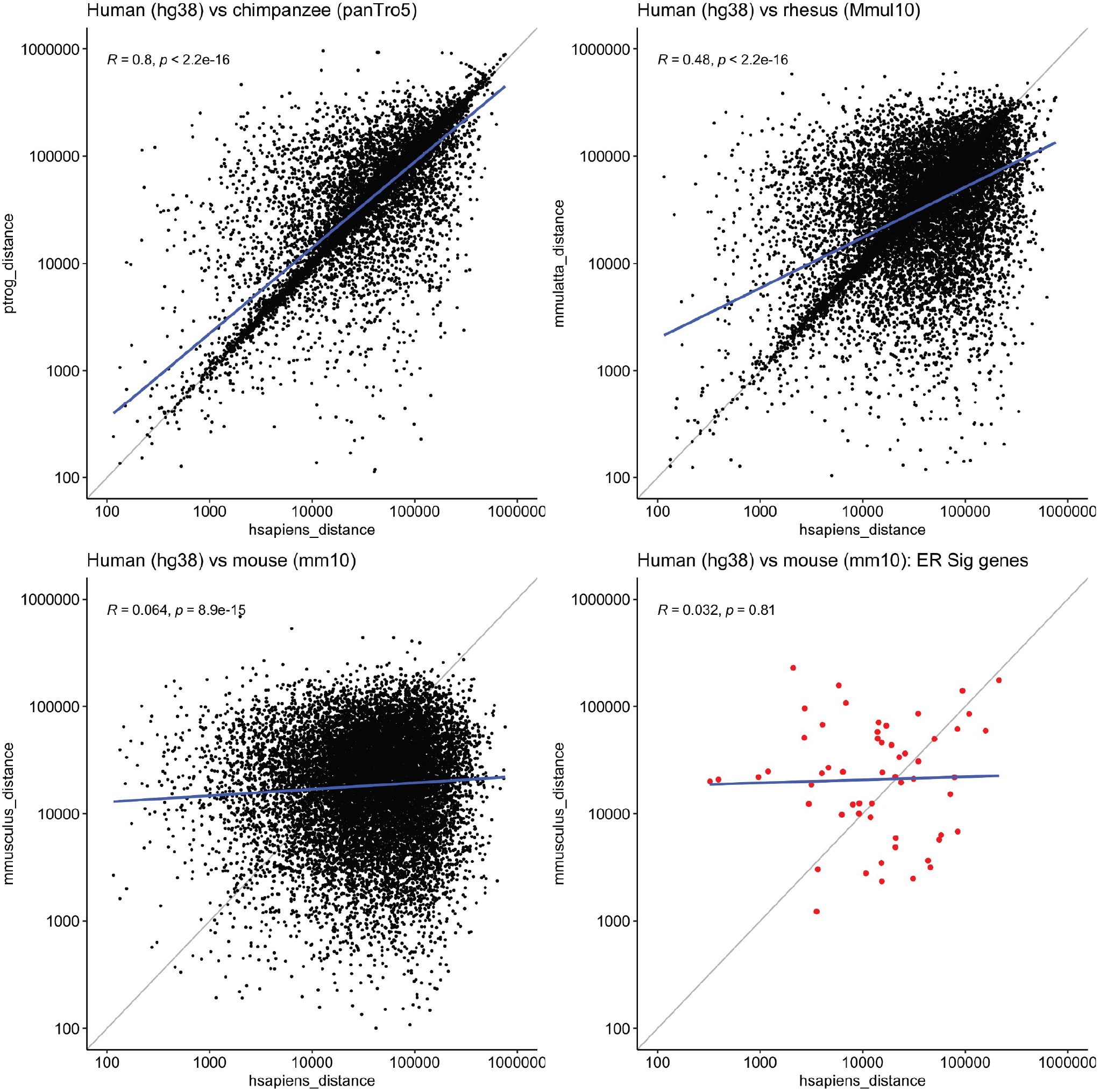
Additional details regarding conservation of relationships between GGAA microsatellites and gene promoters across species. Plots showing distance from transcriptional start sites (TSS) of human genes to the nearest GGAA repeat (3xGGAA) on the x axis, compared to distance from the orthologous gene TSS to the nearest GGAA repeat in the indicated species on the y axis. Linear best-fit line (blue) and Pearson correlation coefficients are indicated. Panel at lower right shows gene-repeat distances for human vs. mouse for previously defined (Ross 2003) E-R signature genes only.

**Supplementary Figure 5:**
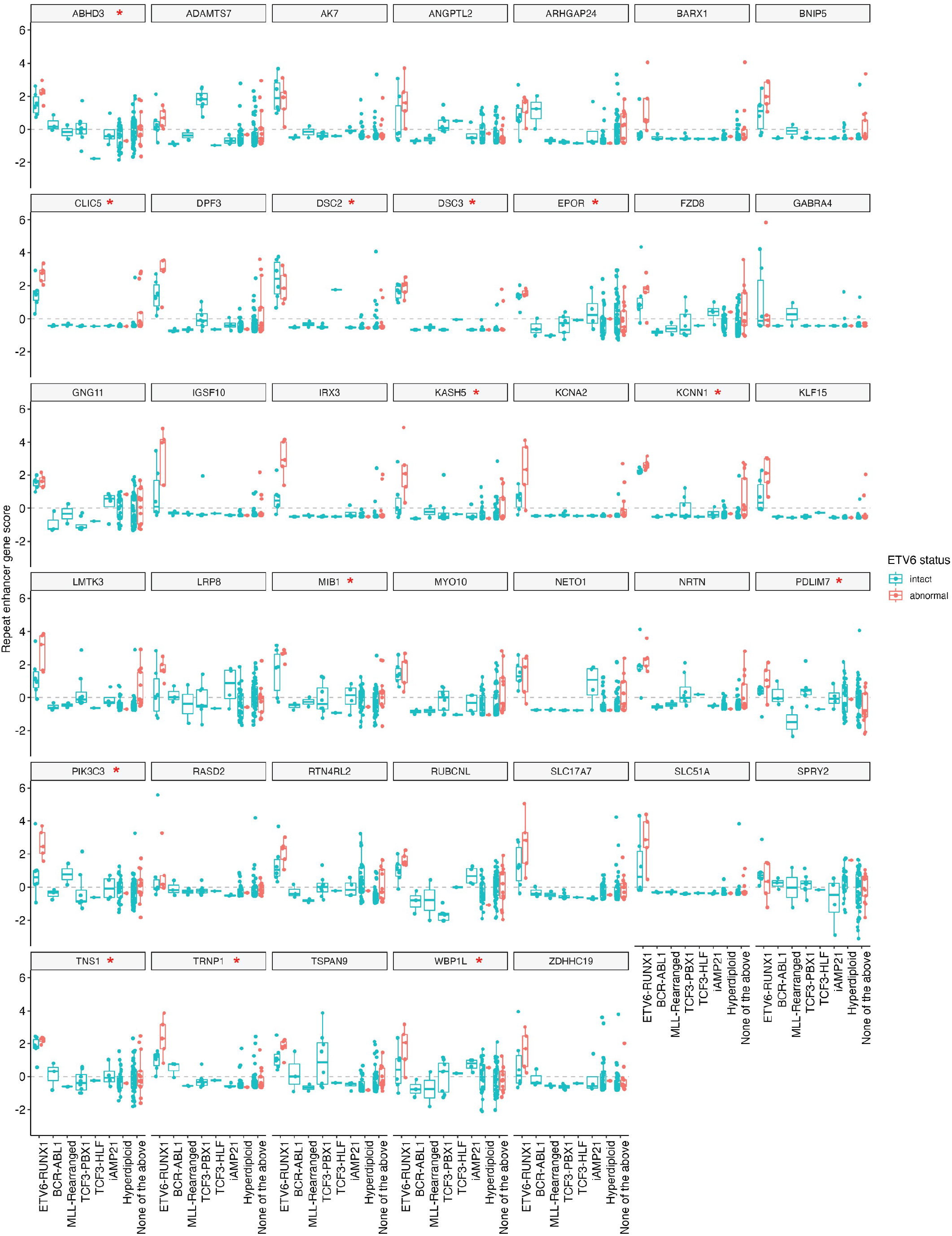
Additional details of ETV6-repressed gene expression in primary B-ALL. RNA-Seq expression Z-scores for the 40 ETV6-repressed genes identified in Reh cells (significant downregulation in ETV6-WT-V5-expressing cells, ETV6-WT-V5 binding site within 200 kbp) that are upregulated in TARGET E-R+ B-ALL samples versus all other samples. Samples are grouped by molecular subtype and ETV6 status as defined in Figure 4B. Gene names marked with a red asterisk are previously reported E-R signature genes (Ross et al, 2003).

**Supplementary Figure 6:**
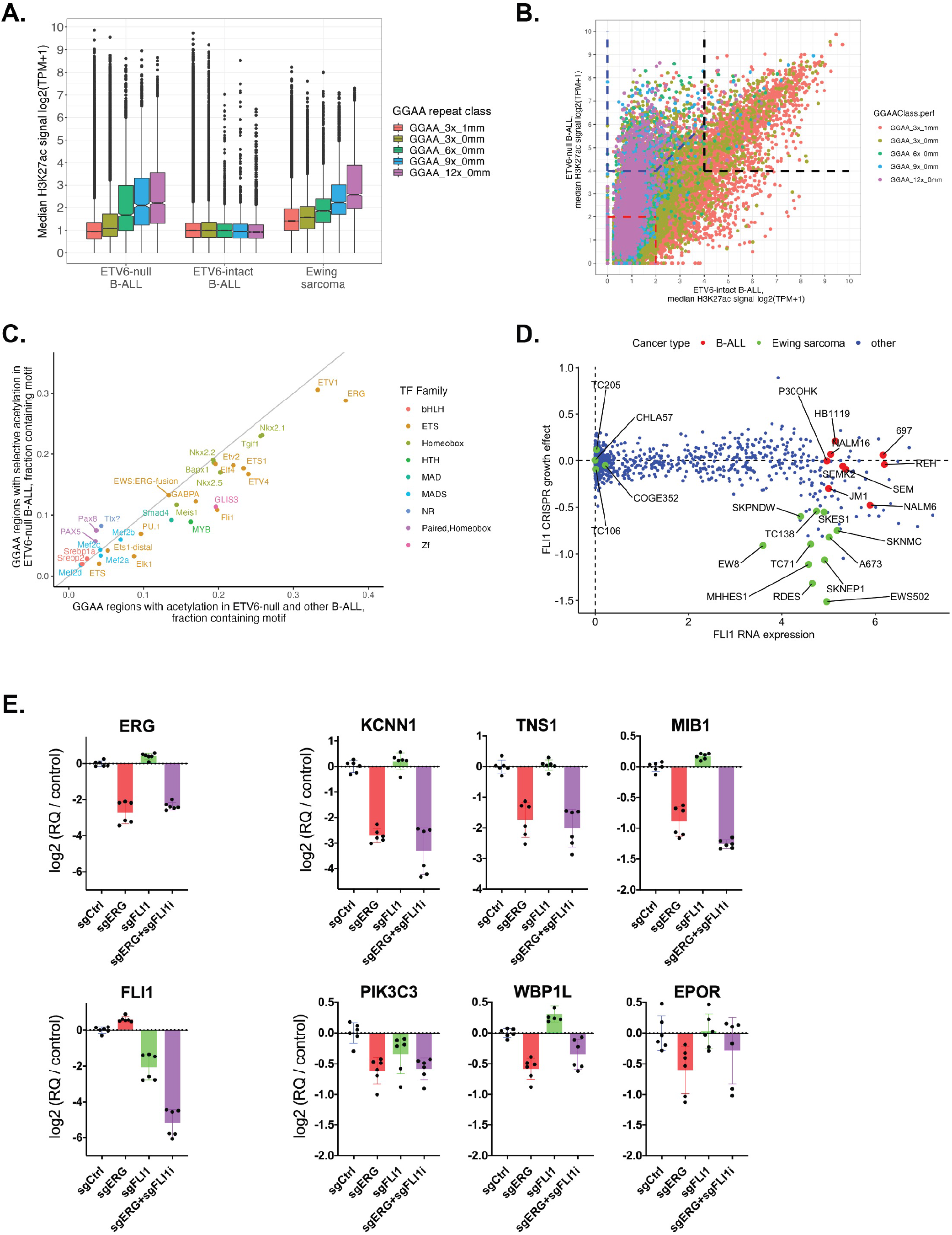
Additional details of GGAA repeat acetylation, motif analysis, and effects of ETS factors in B-ALL. **A.** Comparison of acetylation associated with all hg38 GGAA repeat-containing intervals, classified by length and stringency, for B-ALL cell lines without ETV6 (median of cell lines Reh, AT-1, UoCB6, and MUTZ-5), B-ALL cell lines with intact ETV6 (median of cell lines SUP-B15, NALM-6, SEM, RS4;11, KOPN-8, HAL-01, and MHH-CALL-3), and Ewing sarcoma (median of cell lines A673 and SK-N-MC; data from Riggi et al^41^). **B.** Plot of median H3K27ac signal flanking all hg38 GGAA repeats for B-ALL cell lines without ETV6 and B-ALL cell lines with intact ETV6 (groups defined as in A). Repeats are color-coded by best GGAA repeat class. Dotted lines show thresholds used to define repeats with acetylation specific to E-R+ B-ALL (blue), acetylation in both E-R+ and B-ALL with intact ETV6 (black), and non-acetylated (red) for motif analyses in Figure 6A and Suppl. figure 6C. **C.** Fraction of hg38 GGAA repeat-centered intervals (+/- 100 bp) containing the indicated motif (HOMER motif library v. 4.10) for intervals with E-R-specific acetylation or acetylation in both E-R+ and ETV6-intact B-ALL, as defined in (**B**). Shown are all motifs with enrichment −log(pval) > 8 for the E-R+ intervals versus non-acetylated intervals. **D.** DepMap data showing FLI1 expression and CRISPR knockout growth effect for B-ALL cell lines vs other cancer types. **E.** Differences in transcript levels in Reh cells expressing promoter-targeting sgRNA against *ERG* and / or *FLI1* versus non-targeting sgRNA, 72 hours after doxycycline induction (qRT-PCR, normalized to *EEF1A*). Data are from two separate experiments with three PCR replicates each. Data for sgCtrl and sgErg are the same as shown in Figure 6E.

## Supplementary tables (Excel files)

Table S1: Oligonucleotide sequences and cell line information.

A – Oligonucleotides
B – Cell lines

Table S2: Known and de novo motif analysis for distal acetylation clusters

A – HOMER known motif analysis
B – HOMER *de novo* motif analysis

Table S3: Peak calls and transcription factor enrichment statistics for TF ChIP-Seq

Table S4: ETV6-regulated gene / ETV6 binding site linkages

## References

1. Ross, M. E. et al. Classification of pediatric acute lymphoblastic leukemia by gene expression profiling. Blood 102, 2951–9 (2003).

2. Gu, Z. et al. PAX5-driven subtypes of B-progenitor acute lymphoblastic leukemia. Nat. Genet. 51, 296–307 (2019).

3. Lilljebjörn, H. & Fioretos, T. New oncogenic subtypes in pediatric B-cell precursor acute lymphoblastic leukemia. Blood 130, 1395–1401 (2017).

4. WHO Classification of Tumors of Haematopoietic and Lymphoid Tissues. (World Health Organization, 2017).

5. Lopez, R. G. et al. TEL is a sequence-specific transcriptional repressor. J. Biol. Chem. 274, 30132–30138 (1999).

6. Romana, S. et al. The t(12;21) of acute lymphoblastic leukemia results in a tel-AML1 gene fusion. Blood 85, 3662–3670 (1995).

7. Golub, T. R. et al. Fusion of the TEL gene on 12p13 to the AML1 gene on 21q22 in acute lymphoblastic leukemia. Proc. Natl. Acad. Sci. 92, 4917–4921 (1995).

8. Raynaud, S. et al. The 12;21 translocation involving TEL and deletion of the other TEL allele: two frequently associated alterations found in childhood acute lymphoblastic leukemia. Blood 87, 2891–9 (1996).

9. Kim, D. H. et al. TEL-AML1 translocations with TEL and CDKN2 inactivation in acute lymphoblastic leukemia cell lines. Blood 88, 785–94 (1996).

10. Stegmaier, K. et al. Frequent loss of heterozygosity at the TEL gene locus in acute lymphoblastic leukemia of childhood. Blood 86, 38–44 (1995).

11. Lilljebjörn, H. et al. Identification of ETV6-RUNX1-like and DUX4-rearranged subtypes in paediatric B-cell precursor acute lymphoblastic leukaemia. Nat. Commun. 7, 11790 (2016).

12. Zhang, M. Y. et al. Germline ETV6 mutations in familial thrombocytopenia and hematologic malignancy. Nat. Genet. 47, 180–185 (2015).

13. Nishii, R. et al. Molecular basis of ETV6-mediated predisposition to childhood acute lymphoblastic leukemia. Blood 137, 364–373 (2021).

14. Diedrich, J. D. et al. Profiling chromatin accessibility in pediatric acute lymphoblastic leukemia identifies subtype-specific chromatin landscapes and gene regulatory networks. Leukemia (2021) doi:10.1038/s41375-021-01209-1.

15. 15. Blueprint Epigenome Consortium. http://dcc.blueprint-epigenome.eu/#/datasets.

16. Niebuhr, B. et al. Runx1 is essential at two stages of early murine B-cell development. Blood 122, 413–423 (2013).

17. Linka, Y. et al. The impact of TEL-AML1 (ETV6-RUNX1) expression in precursor B cells and implications for leukaemia using three different genome-wide screening methods. Blood Cancer J. 3, 1–9 (2013).

18. Teppo, S. et al. Genome-wide repression of eRNA and target gene loci by the ETV6-RUNX1 fusion in acute leukemia. Genome Res. 26, 1468–1477 (2016).

19. Wei, G. H. et al. Genome-wide analysis of ETS-family DNA-binding in vitro and in vivo. EMBO J. 29, 2147–2160 (2010).

20. Hollenhorst, P. C., McIntosh, L. P. & Graves, B. J. Genomic and biochemical insights into the specificity of ETS transcription factors. Annu. Rev. Biochem. 80, 437–71 (2011).

21. Therapeutically Applicable Research to Generate Effective Treatments (TARGET) Program, National Cancer Institute. https://ocg.cancer.gov/programs/target/projects/acute-lymphoblastic-leukemia.

22. Inthal, A. et al. Role of the erythropoietin receptor in ETV6/RUNX1-positive acute lymphoblastic leukemia. Clin. Cancer Res. 14, 7196–7204 (2008).

23. Torrano, V., Procter, J., Cardus, P., Greaves, M. & Ford, A. M. ETV6-RUNX1 promotes survival of early B lineage progenitor cells via a dysregulated erythropoietin receptor. Blood 118, 4910–4918 (2011).

24. Polak, R. et al. Autophagy inhibition as a potential future targeted therapy for ETV6-RUNX1-driven B-cell precursor acute lymphoblastic leukemia. Haematologica 104, 738– 748 (2019).

25. Shi, J. et al. Role of SWI/SNF in acute leukemia maintenance and enhancer-mediated Myc regulation. Genes Dev. 27, 2648–62 (2013).

26. Herranz, D. et al. A NOTCH1-driven MYC enhancer promotes T cell development, transformation and acute lymphoblastic leukemia. Nat. Med. 20, 1130–1137 (2014).

27. Bunting, K. L. et al. Multi-tiered Reorganization of the Genome during B Cell Affinity Maturation Anchored by a Germinal Center-Specific Locus Control Region. Immunity 45, 497–512 (2016).

28. Corces, M. R. et al. The chromatin accessibility landscape of primary human cancers. Science 362, (2018).

29. Gunji, H. et al. TEL/AML1 shows dominant-negative effects over TEL as well as AML1. Biochem. Biophys. Res. Commun. 322, 623–630 (2004).

30. Sasaki, K. et al. Functional analysis of a dominant-negative ΔeTS TEL/ETV6 isoform. Biochem. Biophys. Res. Commun. 317, 1128–1137 (2004).

31. De Braekeleer, E. et al. ETV6 fusion genes in hematological malignancies: A review. Leuk. Res. 36, 945–961 (2012).

32. Biswas, A., Rajesh, Y., Mitra, P. & Mandal, M. ETV6 gene aberrations in non-haematological malignancies: A review highlighting ETV6 associated fusion genes in solid tumors. Biochim. Biophys. Acta -Rev. Cancer 1874, 188389 (2020).

33. Suto, Y., Sato, Y., Smith, S. D., Rowley, J. D. & Bohlander, S. K. A t(6;12)(q23;p13) results in the fusion of ETV6 to a novel gene, STL, in a B-cell ALL cell line. Genes Chromosom. Cancer 18, 254–268 (1997).

34. Golub, T. R. et al. Oligomerization of the ABL tyrosine kinase by the Ets protein TEL in human leukemia. Mol. Cell. Biol. 16, 4107–4116 (1996).

35. Yagasaki, F. et al. Fusion of TEL/ETV6 to a novel ACS2 in myelodysplastic syndrome and acute myelogenous leukemia with t(5;12)(q31;p13). Genes Chromosom. Cancer 26, 192–202 (1999).

36. Potter, M. D., Buijs, A., Kreider, B., Van Rompaey, L. & Grosveld, G. C. Identification and characterization of a new human ETS-family transcription factor, TEL2, that is expressed in hematopoietic tissues and can associate with TEL1/ETV6. Blood 95, 3341–3348 (2000).

37. Kim, C. A. et al. Polymerization of the SAM domain of TEL in leukemogenesis and transcriptional repression. EMBO J. 20, 4173–82 (2001).

38. Green, S. M., Coyne, H. J., McIntosh, L. P. & Graves, B. J. DNA Binding by the ETS Protein TEL (ETV6) Is Regulated by Autoinhibition and Self-association. J. Biol. Chem. 285, 18496–18504 (2010).

39. Mackereth, C. D. et al. Diversity in structure and function of the Ets family PNT domains. J. Mol. Biol. 342, 1249–1264 (2004).

40. Gangwal, K. et al. Microsatellites as EWS/FLI response elements in Ewing’s sarcoma. Proc. Natl. Acad. Sci. U. S. A. 105, 10149–54 (2008).

41. Riggi, N. et al. EWS-FLI1 Utilizes Divergent Chromatin Remodeling Mechanisms to Directly Activate or Repress Enhancer Elements in Ewing Sarcoma. Cancer Cell 26, 668– 681 (2014).

42. Tsuzuki, S., Seto, M., Greaves, M. & Enver, T. Modeling first-hit functions of the t(12;21) TEL-AML1 translocation in mice. Proc. Natl. Acad. Sci. 101, 8443–8448 (2004).

43. Fischer, M. et al. Defining the oncogenic function of the TEL/AML1 (ETV6/RUNX1) fusion protein in a mouse model. Oncogene 24, 7579–7591 (2005).

44. Schindler, J. W. et al. TEL-AML1 Corrupts Hematopoietic Stem Cells to Persist in the Bone Marrow and Initiate Leukemia. Cell Stem Cell 5, 43–53 (2009).

45. Rodríguez-Hernández, G. et al. The Second Oncogenic Hit Determines the Cell Fate of ETV6-RUNX1 Positive Leukemia. Front. Cell Dev. Biol. 9, 1–15 (2021).

46. van der Weyden, L. et al. Modeling the evolution of ETV6-RUNX1–induced B-cell precursor acute lymphoblastic leukemia in mice. Blood 118, 1041–1051 (2011).

47. Zhang, L. Q. et al. Establishment of cell lines from B-cell precursor acute lymphoblastic leukemia. Leukemia 7, 1865–74 (1993).

48. Fears, S., Chakrabarti, S. R., Nucifora, G. & Rowley, J. D. Differential expression of TCL1 during pre-B-cell acute lymphoblastic leukemia progression. Cancer Genet. Cytogenet. 135, 110–9 (2002).

49. Naumovski, L. et al. Philadelphia chromosome-positive acute lymphoblastic leukemia cell lines without classical breakpoint cluster region rearrangement. Cancer Res. 48, 2876– 2879 (1988).

50. Riester, M. et al. PureCN: copy number calling and SNV classification using targeted short read sequencing. Source Code Biol. Med. 11, 13 (2016).

51. Huang, C. et al. Proteogenomic insights into the biology and treatment of HPV-negative head and neck squamous cell carcinoma. Cancer Cell 39, 361–379.e16 (2021).

52. Ryan, R. J. H. et al. Detection of Enhancer-Associated Rearrangements Reveals Mechanisms of Oncogene Dysregulation in B-cell Lymphoma. Cancer Discov. 5, 1058– 71 (2015).

53. Ryan, R. J. H. et al. A B Cell Regulome Links Notch to Downstream Oncogenic Pathways in Small B Cell Lymphomas. Cell Rep. 21, 784–797 (2017).

54. Parolia, A. et al. Distinct structural classes of activating FOXA1 alterations in advanced prostate cancer. Nature 571, 413–418 (2019).

55. Kondili, M. et al. UROPA: A tool for Universal RObust Peak Annotation. Sci. Rep. 7, 1– 12 (2017).

56. McKenna, A. & Shendure, J. FlashFry: A fast and flexible tool for large-scale CRISPR target design. BMC Biol. 16, 4–9 (2018).

57. Sanson, K. R. et al. Optimized libraries for CRISPR-Cas9 genetic screens with multiple modalities. Nat. Commun. 9, 1–15 (2018).

